# Identifiability analysis for stochastic differential equation models in systems biology

**DOI:** 10.1101/2020.08.10.245233

**Authors:** Alexander P Browning, David J Warne, Kevin Burrage, Ruth E Baker, Matthew J Simpson

## Abstract

Mathematical models are routinely calibrated to experimental data, with goals ranging from building predictive models to quantifying parameters that cannot be measured. Whether or not reliable parameter estimates are obtainable from the available data can easily be overlooked. Such issues of *parameter identifiability* have important ramifications for both the predictive power of a model, and the mechanistic insight that can be obtained. Identifiability analysis is well-established for deterministic, ordinary differential equation (ODE) models, but there are no commonly-adopted methods for analysing identifiability in stochastic models. We provide an accessible introduction to identifiability analysis and demonstrate how existing ideas for analysis of ODE models can be applied to stochastic differential equation (SDE) models through four practical case studies. To assess *structural identifiability*, we study ODEs that describe the statistical moments of the stochastic process using open-source software tools. Using practically-motivated synthetic data and Markov-chain Monte Carlo (MCMC) methods, we assess parameter identifiability in the context of available data. Our analysis shows that SDE models can often extract more information about parameters than deterministic descriptions. All code used to perform the analysis is available on Github.

## 1 Introduction

Stochastic mathematical models are rapidly becoming an essential tool for interpreting biological phenomena [1–7]. These models are necessitated, in part, by increasing experimental interest in capturing finer-scale, time-series observations [8–12] as well as spatial information [13–18] rather than coarse-scale deterministic trends (figure 1). As computational inference techniques for stochastic models have improved [19–23], a fundamental question that often remains overlooked is whether or not model parameters can be confidently estimated from the available data. Drug development, for example, often relies on the quantification of cell growth rates from a *proliferation assay* (figure 1*a–d*) [24]. If a mean-field model is applied to interpret the most frequently reported observation—cell count data—only the *net* growth rate is identifiable, not the proliferation and death rates [25,26]. Establishing the *identifiability* of model parameters is critical as predictions, and parameter estimates, from a non-identifiable model may be unreliable [27–30], with further analysis required to quantify prediction uncertainty in non-identifiable models [31–33]. Identifiability should always, therefore, be established before parameter estimation is attempted. Such identifiability analysis is well-established for deterministic ordinary differential equation (ODE) models [28,34–41], but there is a scarcity of methods available for the stochastic models that are becoming increasingly important.

**Figure 1.**
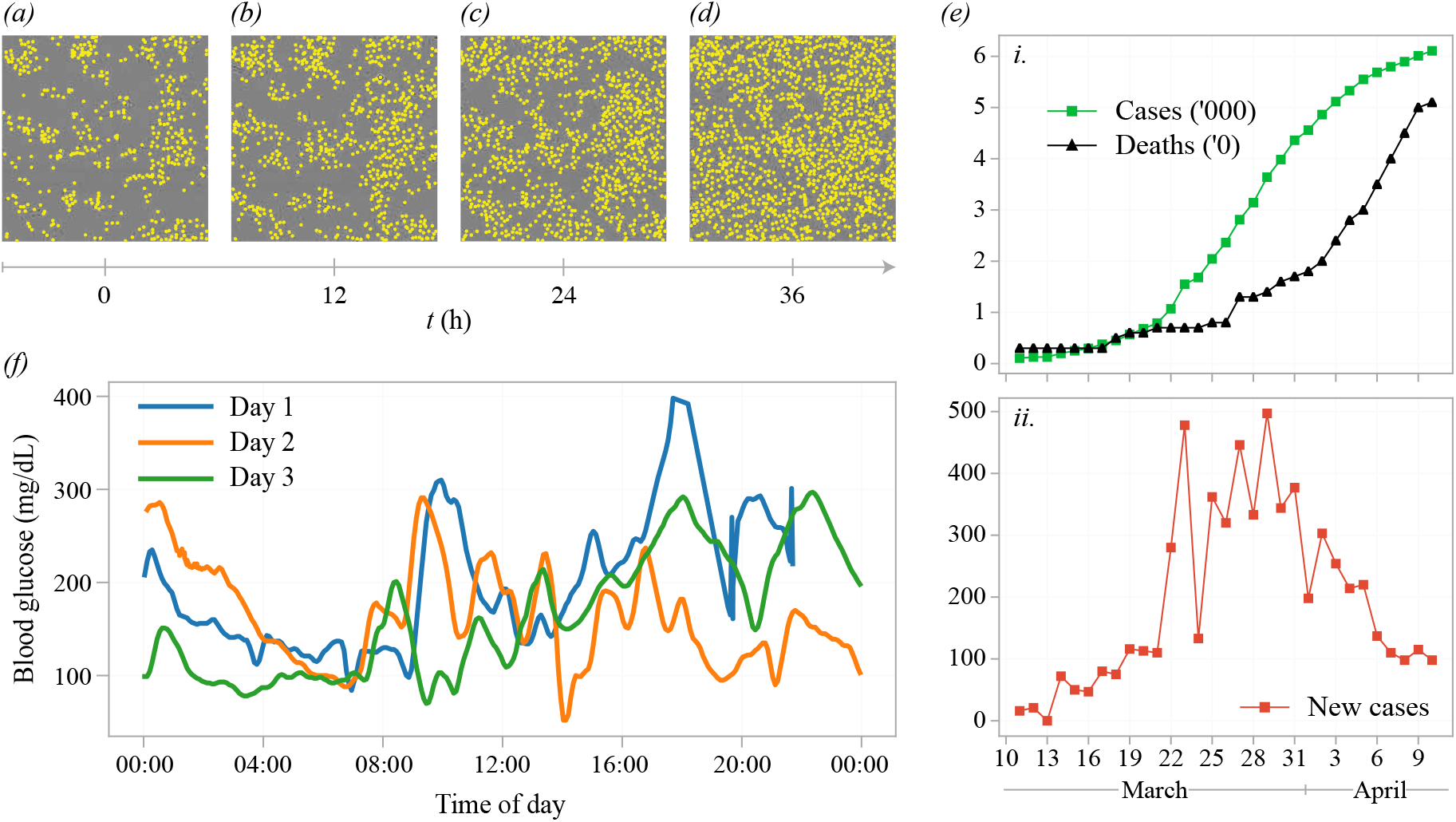
(*a–d*) Cell proliferation and death observed *in vitro* over 36 hours in a proliferation assay [68]. Each snapshot has a field-of-view of 1440 × 1440 μm and the location of each cell is indicated with a yellow marker. (*e*) Data from the early stages of the coronavirus pandemic comprising the observed number of (*i*) infected individuals, deaths, and (*ii*) daily new case count in Australia [69]. (*f*) Continuous glucose monitoring data from a single individual over three consecutive days [70].

Stochasticity is fundamental to many processes [2,42–48]. For example, diabetic patients rely on the rapid interpretation of highly volatile blood glucose measurements to determine insulin input (figure 1*f*) [49,50]. Data from the COVID-19 pandemic [1] is also volatile (figure 1*e*), and inferences of epidemic data must often be drawn from a single, stochastic, time-series. Finally, for systems at equilibrium in the mean-field, such as ion-channel data, models that account for system noise are required to establish parameters [51,52]. Stochastic differential equation (SDE) models of the Itô form are widely applied in systems biology to describe stochastic phenomena [53–56]. While many stochastic systems can be simulated exactly using discrete Markov models, SDE approximations offer a significant computational advantage. In addition, the use of reflected SDEs [57] can guarantee good agreement with their discrete counterparts at boundaries [57,58]. SDE models can describe intrinsic noise in, for example, gene expression [2,9,23] or a bio-chemical reaction network [59]; extrinsic noise describing volatility in the environment [48,53,60,61]; and model approximations and unknown effects in so-called *grey-box* models [62,63]. Explicitly modelling this variability in biological systems can often capture more information about a process than a deterministic model is able to [64–67]. Further, SDE models can account for the correlations inherent to time-series data and account for noise that might otherwise obscure parameters. Our results demonstrate how to establish parameter identifiability for SDE models that encode information about the intrinsic noise of the process [64]. We focus on SDE state-space models that can be formulated through the chemical Langevin equation (CLE), although our intention is to provide analysis applicable to any SDE of the Itô form. The use of such formulations have a long and extensive history of use.

A prerequisite for parameter estimation is that model parameters be *structurally identifiable* [28,34–36,71–73]. Structural identifiability refers to the question of whether a parameter can be identified given an infinite amount of noise-free data. A state-space model is said to be structurally identifiable if distinct values of the parameters imply distinct observed model outputs (or in the case of a stochastic model, distinct observed output distributions [74]), and vice versa [75–77]. Techniques such as differential algebra [38, 78, 79] and transfer function approaches [34, 35] can establish structural identifiability in ODE models. These approaches are also used to establish identifiable relationships between parameters [35,80]—for example, the net growth rate in a proliferation assay—which can aid model design and model reduction [80–82]. Many of these techniques have accessible implementations in symbolic computation packages [38,83–86], meaning structural identifiability analysis does not require a detailed understanding of the, often complex, underlying mathematical analysis [38].

When experimental data is considered, a more useful question is that of *practical identifiability* or *estimability* [28,77,83,87]. That is, can parameters in the model be accurately estimated given a finite amount of noisy experimental data? This kind of analysis is routinely used in the field of experimental design to assess the nature of data required to adequately identify biophysical parameters [29,51,55,88–90]. Practical identifiability is established in conjunction with an inference technique, such as profile or maximum likelihood [90–93] or Markov-chain Monte-Carlo (MCMC) [29, 51]. These techniques provide information about the flatness (or otherwise) of the likelihood function or, in the Bayesian case, the posterior distribution, that describes knowledge about the parameters after the experimental data is taken into consideration. For deterministic and simple stochastic models, this information can be obtained directly from the Fisher information matrix [93]. A model parameter is classified as practically non-identifiable if it cannot be established uniquely within a reasonable level of confidence [28,29]. Compared with structural identifiability, which is a property of the model, practical identifiability is more nuanced and additionally dependent upon prior knowledge; the experimental data; and consequentially, the experiment itself [29,83]. For example, should the model and data provide no more information about a parameter than that already established in previous studies, the parameter may be classified as practically non-identifiable from the data and model in question. For this reason, we take a Bayesian approach to parameter estimation and encode existing knowledge about the parameters in a *prior distribution*. This question of practical identifiability has not yet been demonstrated for SDE models in systems biology.

Computational inference for stochastic models is a significant challenge [22]. Unlike approaches to parameter estimation for deterministic models, the likelihood function for a realistic stochastic model is, generally, intractable [22]. Techniques based on approximations, such as a linear-noise approximation [93] or approximate Bayesian computation [20,94–98], are available for SDEs but are, naturally, approximations. Pseudo-marginal methods [99,100], developed relatively recently, are computationally costly, but provide an unbiased estimate of the true likelihood function for partially observed time-series described by non-linear stochastic models. In this review, we utilise a pseudo-marginal MCMC approach, where we estimate the likelihood with a particle filter, which we refer to as particle MCMC [101,102]. There are many excellent reviews of inference for stochastic models in systems biology [20,22,102,103], so we do not focus on the details our out implementation here. Despite the established importance of identifiability, it is all too common in parts of the inference literature to draw the standard assumption that the model parameters are identifiable: we note that all the aforementioned review articles make no mention of identifiability. The computational cost of inference for stochastic models, in itself, motivates us to consider identifiability. For example, identifiability can guide model selection: if both a deterministic and stochastic description of a process are practically non-identifiable, the cheaper deterministic model may, in some cases, be adequate for parameter estimation. Where structural non-identifiability is detected, practical non-identifiability necessarily follows and does not need to be established separately.

The focus of this review is to provide an accessible guide to establishing identifiability in SDE models in biology. To do this, we analyse identifiability in SDE descriptions of four case study models, shown in figure 2. The simplest model we consider is a birth-death process (figure 2*a*) that is routinely used to describe cell proliferation and death in a range of *in vitro* and *in vivo* biological systems, such as that shown in figure 1*a*. We demonstrate that, from cell count-data, the cell proliferation and death rates are structurally non-identifiable for a routinely employed ODE model, but can be identified for an SDE model. Next, we consider two multi-state models where only partial observations of the system are available. First, a two-pool model (figure 2*b*) that can describe, for example, the decay of human cholesterol whilst it transfers between two organs [35,104]. We assume that data from the two-pool model comprises several time-series observations of the substance concentration in a single pool. Secondly, an epidemic model (figure 2*c*) [105–107] describes individuals infected due to interactions between susceptible and infectious individuals. We model a testing procedure such that unknown proportions of the number of infectious and recovered individuals are observed, and inferences are drawn from a single time-series. The last model we consider is a non-linear SDE model for insulin regulation by *β*-cells (figure 2*d*) [108,109]. This type of model can describe the volatility associated with data from a continuous glucose monitoring device (figure 1*f*) [70]. The equivalent ODE description of the *β*-insulin-glucose circuit is not structurally or practically identifiable [110], and we demonstrate how the analysis for the ODE description can inform a parameter transformation to aid identifiability analysis for the SDE model.

**Figure 2.**
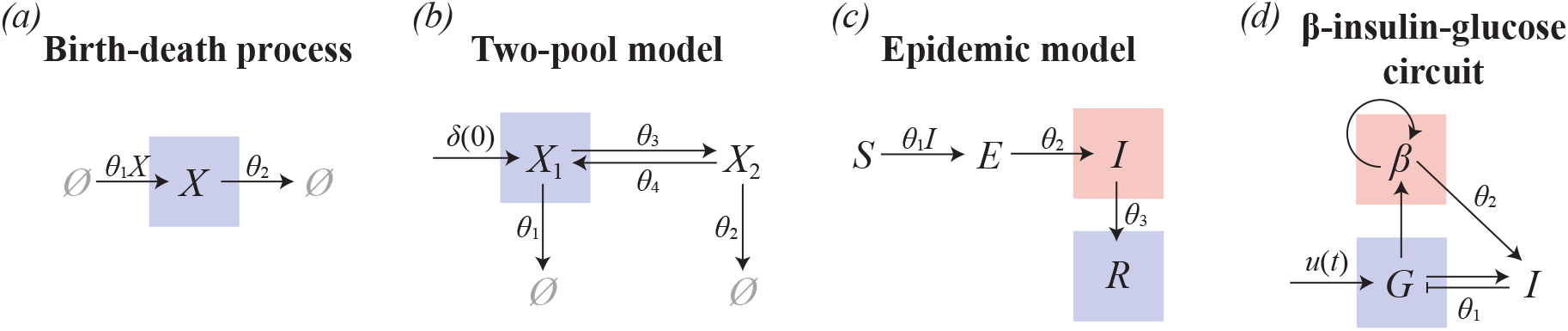
We demonstrate identifiability in an SDE CLE description of four models: (*a*) a birth-death process; (b) a two-pool model; (c) an epidemic model; and (d) a *β*-insulin-glucose circuit. The coloured boxes indicate the observed quantity, which is coupled to a noisy observation process.

We demonstrate two main approaches to assess identifiability in SDE models. First, we assess structural identifiability through a surrogate model, taken to be a system of ODEs that describe the time-evolution of the statistical moments of the SDE [111–115]. This allows us to apply the established open-source structural identifiability software package `DAISY` (written for the freeware `REDUCE` software) to the SDE models through the moment equations. We repeat this analysis in the more recent open-source software package `GenSSI2` [116,117], written for `MATLAB`, which can be more efficient for non-linear systems. We interpret these results as a proxy for identifiability of the SDE model itself. While this approach is not always conclusive, it can provide a rapid preliminary screening tool and allows direct comparison of identifiability for an SDE model, which contains information about the mean, variance and higher moments; to identifiability for a corresponding ODE model that is typically assumed to describe an approximation of the mean. We only apply this approach where an exact system of moment equations can be derived, which occurs when the reaction rates are polynomial. For more complex stochastic models containing terms such as Hill functions, as found in the *β*-insulin-glucose circuit model, an exact system of moment equations cannot be derived, we do not apply the moment dynamics approach in this case. We assess practical identifiability for all models using MCMC [29,51], first demonstrating how practical identifiability can be cheaply established from a naïve proposal kernel. To compute credible intervals for each parameter, and visualise potential correlations between parameters, we produce results using a tuned proposal kernel where we can be more certain of convergence.

The outline of this review is as follows. In Section 2, we establish the types of SDE models and observation processes that we consider, and then outline the techniques used to generate synthetic data. Following this, in Section 2.2, we summarise moment closure techniques for SDEs and describe how we implement the software tools `DAISY` and `GenSSI` to assess for structural identifiability. Next, in Section 2.3, we provide a brief overview of our implementation of the particle MCMC algorithm. Full details of particle MCMC for SDE models can be found in the existing literature [102,103] and as supporting material. In Section 3, we use these tools to assess identifiability using an SDE description of four models. In Sections 4 and 5, we discuss our results and provide an outlook on the future of identifiability for stochastic models in biology. To aid in the accessibility of the techniques we review, we provide our MCMC code in the form of a module^1^ for the open-source, high-performance `Julia` programming language [118].

## 2 Mathematical techniques

In this section, we outline the mathematical and statistical techniques we use to perform identifiability analysis. Full details of all algorithms used are provided as supporting material.

### 2.1 Stochastic models in biology

We consider Itô SDE state space models of the form

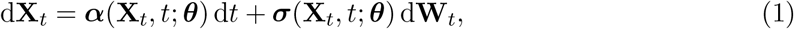

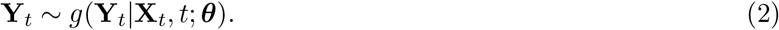

Here, the state is described by 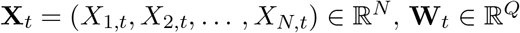 is a *Q*-dimensional Wiener process with independent components; ***α***(·) maps to an *N*-dimensional vector; and ***σ***(·) maps to an *N × Q* matrix. The observables, 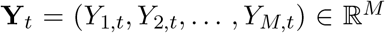, are connected to the state variables according to an observation process with probability density function *g*(**Y**_*t*_|**X**_*t*_, *t*; ***θ***). We consider several forms of observation function, including partial observations of the state with both additive and multiplicative Gaussian noise with unknown variance 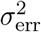. In equations (1) and (2), ***θ*** is a vector of unknown parameters to be determined through inference. In this review, all variables and parameters are dimensionless.

The focus of this review is on Itô SDE models that are formulated through the CLE description of a system of bio-chemical reactions [59,119,120]. Therefore, additional information about rate parameters is encoded in the noise of the process. The first three models we consider (figure 2*a–c*) can be expressed directly as a network of reactions. As the *β*-insulin-glucose circuit model (figure 2*d*) involves state variables modelled as concentrations, not individual counts, we derive a stochastic description from the CLE but scale the noise term in proportion to the concentration of each species.

In summary, a bio-chemical reaction network comprises *N* species, *X*_1_, *X*_2_,…, *X_N_*, that interact through *Q* reactions [121–123]. The population of each species is given by 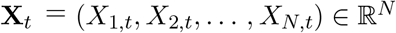. By the law of mass action [59,124], each reaction occurs with a rate described by a *propensity function, a_k_*(**X**_*t*_, *t*; ***θ***), which is equal to the product of the reactants and the rate constant. The net effect of the kth reaction is described by the stoichiometry ***ν**_k_* such that, should reaction *k* occur in [*t, t* + d*t*),

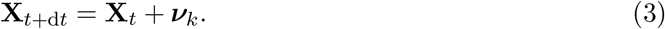

For bio-chemical reaction networks without an explicit time-dependent input, the propensity functions will be independent of *t* and the system can be simulated exactly using an event-driven stochastic simulation algorithm (SSA) [5,124–126]. The principle behind an exact SSA is that reactions can be modelled by an inhomogeneous Poisson process. The time interval between reactions, Δ*t*, is exponentially distributed such that

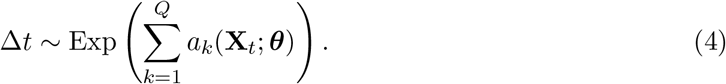

A single reaction occurs at each time-step; the kth reaction occurs with probability proportional to *a_k_*(**X**_*t*_; ***θ***). A typical implementation of the SSA first samples a time-step using equation (4); then samples the next event to occur; and finally updates the state. Full details of our implementation of an SSA are given as supporting material, and the reader is directed to [121] for a comprehensive review of simulation algorithms for bio-chemical reaction networks. We generate synthetic data for the first three models, for which the propensity functions are independent of *t*, using the SSA. In figure 3*a–c* we show 100 realisations of the SSA for the birth-death process, two-pool model and epidemic model, respectively.

**Figure 3.**
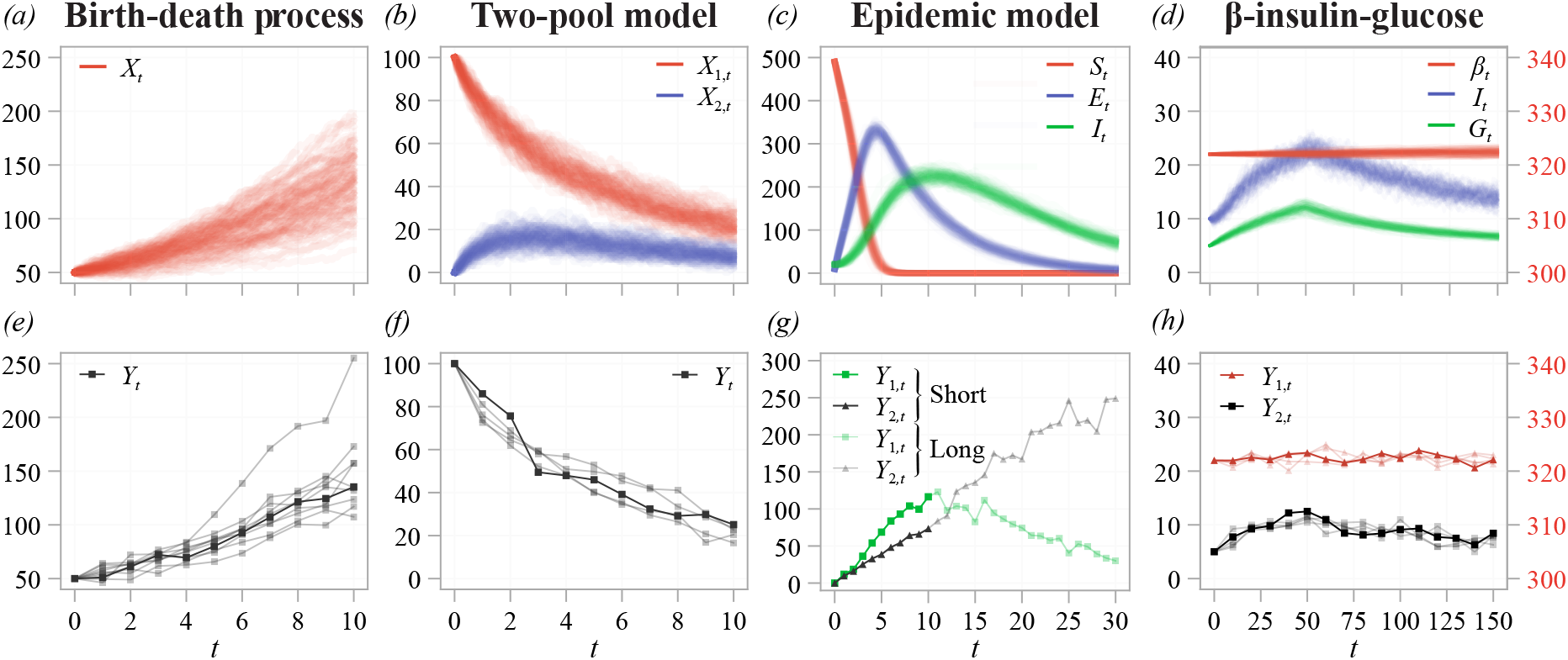
(*a–d*) 100 example realisations of each model, produced using: (*a–c*) the SSA; and, (*d*) the SDE. (*e–h*) Synthetic data used for practical identifiability analysis. Synthetic data comprises noisy observations of the (*e*) full and (*f–h*) partial state. In (*e,f,h*), experimental replicates used simultaneously for parameterisation are shown semi-transparent, with the first replicate fully opaque. For the epidemic model, both short-time (opaque) and long-time (semi-transparent) data are considered separately. In both cases of the epidemic model, an unknown proportion of the number of infected individuals (green), and the recovered individuals (black), is observed. In (*d,h*), the *β* cell concentration, *β_t_* (and the measured concentration *Y*_1,*t*_) is shown on the right axis.

When the population of each species is large and reactions sufficiently frequent, the dynamics of a bio-chemical reaction network can be approximated using the CLE [25,119,121]. Such an approximation is widely applied in systems biology [127, 128], and it is often necessary as the SSA quickly becomes computationally expensive as the populations become large and reactions are frequent enough [129]. The CLE is an Itô SDE of the form

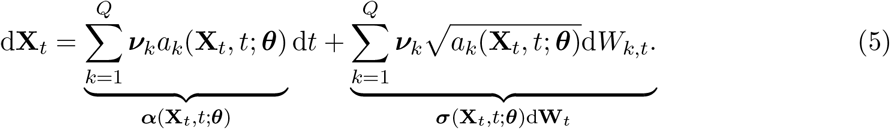

Here **W**_*t*_ = (*W*_1,*t*_, *W*_2,*t*_,…, *W_Q,t_*) is a *Q*-dimensional Wiener process with independent components. In this study, we derive an SDE description for each model using the CLE, and we calibrate this SDE to the synthetic data to approximate the parameters in each model. For the first three models, where data is generated using the SSA, not the SDE, this means that identifiability analysis is conducted in such a way that model misspecification could potentially arise. This pragmatically mirrors experimental data, where any model (including an ODE and SDE description) is an approximation. The forward simulation for each SDE is approximated using the Euler-Maruyama algorithm [130], where we apply reflected SDEs to ensure positivity [57]. Full details of the numerical algorithm are given as supporting material.

### 2.2 Moment dynamics

To enable the application of established methods for structural identifiability analysis to SDE models, we formulate a system of ODEs that describe the statistical moments of the random variable 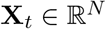. We denote *m*_*i*_1_ *i*_2_…*i_d_*_(*t*) as a raw moment of **X**_*t*_, such that [112–114,120]

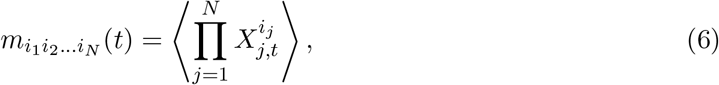

where 〈·〉 indicates the expectation taken with respect to the probability measure of the random variable **X**_*t*_. Here, 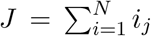 is the *order* of the moment. For example, the first order moments correspond to the mean of each dimension of **X**_*t*_, the second order moments relate to the variances and covariances, and so forth.

We apply the software packages `DAISY` [38] and `GenSSI2` [117] to establish structural identifiability of the resultant system of moment equations. The software package takes a system of ODEs describing the state equations—in our case, the moment equations—in addition to an explicit algebraic relationship between the observables and the state. We, therefore, provide the moments of the observables, **Y**_*t*_, in the noise-free limit, which we denote

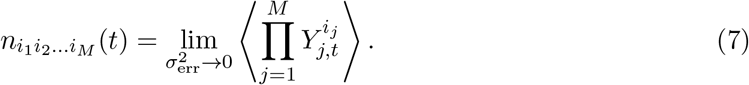

In many cases, the observation distribution, *g*(**Y**_*t*_|**X**_*t*_, *t*; ***θ***), will depend upon the unknown parameters, ***θ***, if, for example, an unknown proportion of the state is observed. This is captured in the structural identifiability analysis as the equations derived for the observed moments, *n*, may depend on ***θ***. We provide well commented input and output obtained using `DAISY` on Github as supporting material.

An expression for the time derivative of each moment can be found using Itô’s lemma (supplementary material). When each component of ***σ***^*T*^***σ*** is an analytic function, which occurs when all the propensity functions in the bio-chemical reaction network are also analytic functions, we obtain [131]

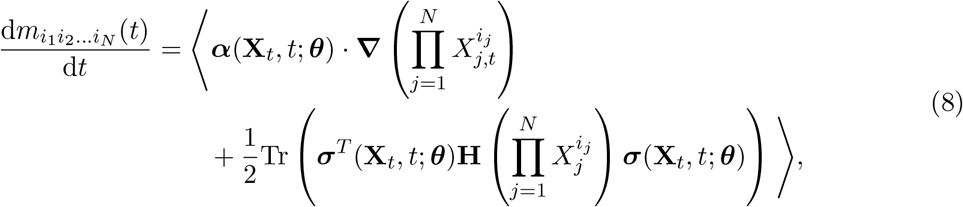

where **H**(·) denotes the *N × N* Hessian matrix of its argument and 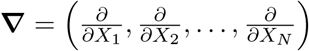. In the case that *N* = 1, equation (8) reduces to

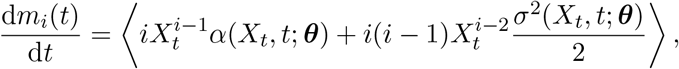

where *α* and *σ* are now scalar functions.

When each component of ***α*** and ***σ**^T^**σ*** are polynomials in **X**_*t*_, the expectation in equation (8) can be carried through to replace powers of **X**_*t*_ with appropriate moments. This, in general, provides an infinite system ODEs that *exactly* describe the time evolution of the moments. In practice, we consider a finite system of moments, up to and including moments of order *J*. We express this now finite system of ODEs as

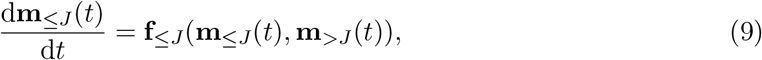

where **m**_≤*J*_(*t*) is a vector containing all the moments up to, and including, order *J*; and **m**_>*J*_(*t*) is a vector containing all moments of order *J* + 1 and above. In the case that **f**_≤*J*_(·) depends only on moments up to order *J*, the system is said to be *closed* at order *J*. That is, the infinite system of equations can be truncated at order *J* and solved directly to obtain an exact solution for the moments. This is the case if ***α*** and ***σ**^T^**σ*** are linear in **X**_*t*_, which occurs in SDEs derived from the CLE if each propensity is linear in **X**_*t*_, as is the case for the first two models we consider (figure 2*a,b*).

For more complicated models, including the epidemic model (figure 2*c*), the system will not, in general, be closed. We must, therefore, apply a *moment closure* approximation to express moments of order higher than *J* in terms of lower order moments [45]. Moment closures typically make an *a priori* assumption about the distribution of the random variable **X**_*t*_. For example, assuming components of **X**_*t*_ are independent or normally distributed is a common approach. In this review, we consider three common moment closures: (1) a mean-field closure [113]; (2) a pairwise closure [113]; and (3) a Gaussian closure [112].

The *mean-field closure* we consider makes the approximation

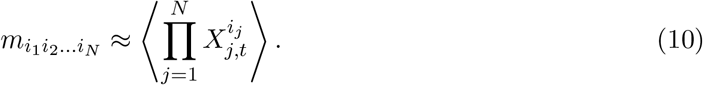

This closure is derived from the assumption that components of **X**_*t*_ are weakly correlated [113] and is also referred to as the *covariance closure* [132]. In the case a closure is drawn at *J* = 1, the mean-field closure often corresponds to an ODE description of the process. For our analysis of the epidemic model, we find that the mean-field closure behaves poorly, suggesting that an assumption that the components of **X**_*t*_ are independent may not be appropriate (Supplementary Material, Figure S1).

While the mean-field closure is commonly drawn at order *J* = 1, it is more common for the *pair-approximation closure* to be applied for second and higher order closures [113]. The pair-approximation closure assumes that a third order moment can be expressed as

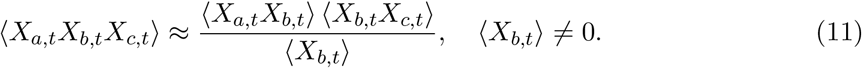

The *Gaussian closure* approximates higher order moments to match those of the normal distribution, and gives a closure in terms of the mean and covariances. Higher order moments can be approximated with [112, 133]

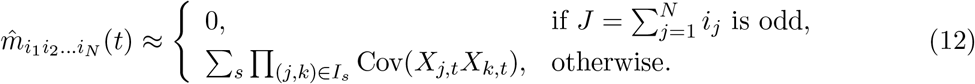

Here, 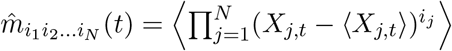 denotes a central moment; Cov(*X_j,t_X_k,t_*) denotes the covariance between *X_j,t_* and *X_k,t_*; and *I_s_* are the sets formed by partitioning the set 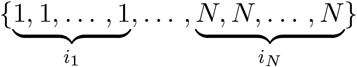 into unordered pairs, where *s* is the number of sets. The raw moments, *m*_*i*1*i*_2_…*i_N_*_(*t*) can then be solved from the expressions for the central moments obtained from equation (12). For a practical example of the Gaussian closure, see [112].

Other choices of moment closure are routinely used in systems biology, such as those based upon a multivariate lognormal distribution [112] or a derivative matching scheme [134]. However, more complex closures add further complexity to the moment equations, which is a significant computational disadvantage for automated assessment of structural identifiability in software packages such as `DAISY` and `GenSSI2`. Furthermore, an *approximate* system of moment equations (which must then also be closed) could be obtained by applying a series expansion approximation, or an approximation similar to the mean-field closure, to systems containing non-polynomial analytic functions; this is the case for the fourth model we consider (figure 2*d*). We do not consider the moment dynamics approach for non-polynomial models in this review.

### 2.3 Inference with MCMC

We take a Bayesian approach to parameter estimation to update our knowledge about the parameters, ***θ***, from a set of observations, 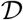, using the likelihood function, 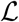, such that [135]

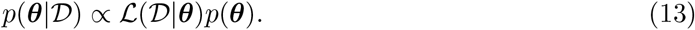

Here, *p*(***θ***) is the *prior distribution*, and represents our knowledge of ***θ*** before consideration of the observations 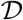. The prior distribution may encode information from, for example, previous experiments, established knowledge, or physical restrictions on the parameters. In the context of practical identifiability, our goal is to significantly increase our understanding of ***θ*** from our prior knowledge. We specify *p*(***θ***) to be a truncated uniform distribution: all parameters within a specified region of realistic parameter values (the support) are considered equally likely [29]. An advantage of a uniform prior in the context of identifiability is that the posterior corresponds to the truncated likelihood function, and, therefore, high density regions of the posterior correspond to regions of high likelihood. Further, should an improper, unbounded uniform prior be considered, the posterior will be directly proportional to the likelihood. Thus, our methodology can also be applied to assess parameter identifiability using a purely likelihood-based approach.

We use an MCMC technique, based on the Metropolis-Hastings algorithm, to sample from the posterior distribution [135–138]. The principle behind MCMC in Bayesian inference is to construct a Markov chain, {***θ**_i_*}_*i*≥0_, with a stationary distribution equal to 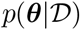. We make a standard choice to initiate the chain from a prior sample, ***θ***_0_ ~ *p*(***θ***). At each iteration of the algorithm, a new state is proposed, ***θ***\* ~ *q*(***θ***\*|***θ**_m_*), where *q* is termed the *proposal kernel*. The proposal is accepted, ***θ***_*m*+1_ **← *θ*\***, with probability

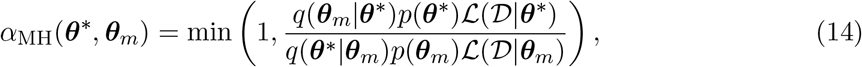

else the proposal is rejected, ***θ***_*m*+1_ **← θ**_*m*_. Full details of our implementation are provided as supporting material. In this review, we use a multivariate normal proposal so that *q*(***θ***_*m*_|***θ***\*) = *q*(***θ***\*|***θ**_m_*). An interpretation of the Metropolis choice of acceptance probability, equation (14), where the proposal is normal and, therefore, symmetric, is that proposals that increase the posterior density are always accepted, whereas proposals that decrease the posterior density are accepted with some reduced probability [29].

We refer to the first set of MCMC chains for each problem as *pilot chains* [139]. The proposal distribution for each pilot chain is set to be a multivariate normal distribution with independent components and variances equal to one-tenth the corresponding prior variance for each parameter, a typical choice. We always produce four pilot chains, each of 10,000 iterations, which we find to be sufficient to indicate identifiability for our models. These pilot chains are then used to *tune* the MCMC proposal kernel [140]. We then produce four *tuned chains*, which can be reliably used to estimate credible intervals and other features of the posterior distribution. The proposal distribution for each tuned chain is chosen to be multivariate normal, with covariance given by [139]

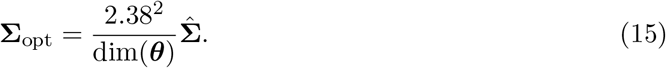

Here, dim(***θ***) is the number of unknown parameters, and 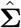 is the covariance matrix for the pooled samples from the four pilot chains (a total of 28,000 samples after 3,000 samples are discarded as burn-in from each pilot chain). To assess convergence, we calculate the commonly used 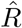 [141] and *n*_eff_ (effective sample size) [135] diagnostics. In summary, 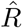 measures the ratio of between-chain and within-chain variance; and *n*_eff_ measures the effective number of independent samples drawn from the posterior. To draw reliable inferences, Gelman *et al*. [135] suggest ensuring that 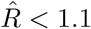. Full details of these convergence statistics are available in [135].

The primary challenge with performing inference for SDE models, with time-series data, is computing the likelihood function. In this review, we consider synthetic data from *E* independent experiments, each with *N_E_* time-series observations. The data are denoted

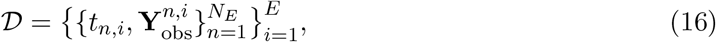

and correspond to the likelihood function

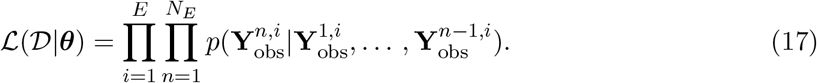

In most cases, the likelihood for noisy time-series data modelled by an SDE will be intractable [102]. This contrasts with data modelled by a deterministic model, which are typically assumed to be independent and normally distributed about the model output [29]. Likelihood free methods, such as ABC [20,97] and pseudo-marginal approaches [100], are routinely used in systems biology to calibrate complex stochastic models to experimental data by approximating equation (17). In this study, we apply a pseudo-marginal approach based on a bootstrap particle filter to approximate the likelihood and calibrate each SDE model to synthetic experimental data [102]. In summary, the bootstrap particle filter approximates equation (17) by

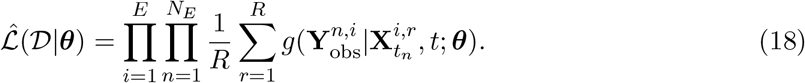

Here, the observation probability density, *g* (equation (2)), is averaged over *R* samples from the SDE, 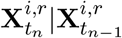 to approximate the likelihood. The bootstrap particle filter then resamples from the set of weighted samples, 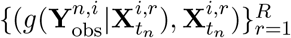, at each time-step to form the starting locations for each SDE sample to sample forward to *t*_*n*+1_. This process is repeated for each independent experiment, and the result is an unbiased Monte Carlo estimate of the likelihood function, 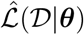, that replaces 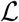 in the Metropolis acceptance probability (equation (14)). Full details of the particle MCMC algorithm, including an implementation for an ODE model used in one case study, are provided as supporting material, and for further information the reader is directed to [102, 103].

## 3 Case studies

Using the moment equations and MCMC, we provide a practical guide for assessing parameter identifiability in SDE CLE models through four case studies. We generate synthetic data for each model using the SSA when the propensity functions are time-independent (the birth-death process, two-pool model and epidemic model), and the corresponding CLE when the propensity functions are time-dependent (the *β*-insulin-glucose circuit). In practice, we would first assess practical identifiability using the experimental data available. However, working with synthetic data provides the means to evaluate the effect of different experiment designs, and observation protocols, on practical identifiability. Our focus is on data comprising partial observations of the process that realistically captures potential experimental data.

### 3.1 Birth-death process

The first model we consider is a birth-death process (figure 2*a*). The birth-death processes can describe, for example, the growth of a well-mixed cell population where individuals proliferate and die according to rates *θ*_1_ and *θ*_2_, respectively. We consider practical identifiability for synthetic data comprising noisy measurements of the cell count at 10 equally spaced times in 10 identically prepared experiments. Such data are typical for *in vitro* cell proliferation experiments [24,142], an example of which is shown in figure 1*a–d*.

#### 3.1.1 Model formulation and moment equations

The birth-death process can be expressed as the bio-chemical reaction network

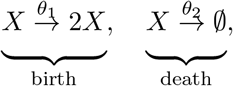

with stoichiometries *ν*_1_ = 1 and *ν*_2_ = −1; and propensities *a*_1_(*X_t_*) = *θ*_1_*X_t_* and *a*_2_(*X_t_*) = *θ*_2_*X_t_*. Here, we denote *X_t_* as the number of individuals in the population. The observed number of individuals, *Y_t_*, is described by the noise model

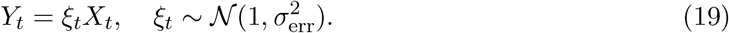

Here, we consider a noise process that scales with the total population, that is, multiplicative Gaussian noise. We show 100 realisations of the SSA for the birth-death process in figure 3*a*, and the synthetic data used for practical identifiability analysis in figure 3*e*. The data are generated using the initial condition *X*_0_ = 50 and target parameter values *θ*_1_ = 0.2, *θ*_2_ = 0.1 and *σ*_err_ = 0.05. Here, *σ*_err_ ≪ 1, which ensures that *Y_t_* remains positive.

The CLE for the birth-death process is

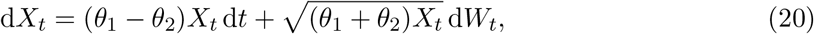

and the first and second order moment equations are

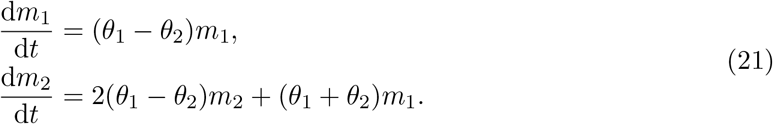

The moment equations for the SDE description of the birth-death model above are identical to the moment equations for the discrete Markov model that we simulate using the SSA [143]. The moments of the observable (in the noise-free limit) are given to second order by *n*_1_ = *m*_1_ and *n*_2_ = *m*_2_. As *α*(·) and *σ*^2^(·) are linear in *X_t_*, the moment equations of the birth-death process are closed at every order and so equations (21) are exact. Further, we note that the common mean-field model for the birth-death process,

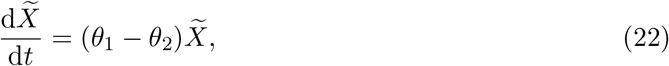

corresponds to the first moment, and describes the average behaviour of *X_t_*. The solution to equation (22) is

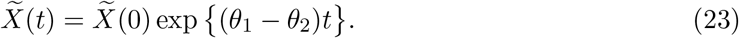

Here, the population, 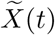, undergoes exponential growth with a net-growth rate of *θ*_1_ − *θ*_2_. Therefore, intuitively, it is not possible to identify *θ*_1_ and *θ*_2_ if only average growth behaviour is observed [25].

#### 3.1.2 Structural identifiability

We first assess structural identifiability of the moment equations in `DAISY` [38]. If only the first moment, *n*_1_, is observed, the system is structurally non-identifiable, meaning the model parameters cannot be uniquely estimated with any amount of data. However, the system becomes structurally identifiable if *n*_2_ is also observed. As the moment equations are closed at every order, and therefore exact, this analysis indicates that the ODE model (equation (22), corresponding to the first moment equation) is structurally non-identifiable, while the SDE model is structurally identifiable.

These structural identifiability results can be intuitively understood through reparameterisation [40]. The first moment equation (or the ODE model) can be re-parameterised with 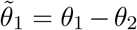 where 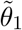 is the sole parameter in the model. Therefore, for a fixed 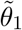, all values on the line 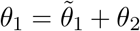 produce indistinguishable behaviour in the first moment, *m*_1_, and hence in the observation, *n*_1_. On the other hand, when re-parameterised the second moment equation contains a second, linearly independent, parameter 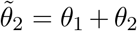. For the birth-death process, the second moment provides enough additional information to uniquely identify both parameters *θ*_1_ and *θ*_2_, provided enough data is available. Thus, the birth-death process is structurally identifiable from the first two moments.

#### 3.1.3 Practical identifiability

We assess practical identifiability of the parameter vector ***θ*** = (*θ*_1_, *θ*_2_, *σ*_err_) for the ODE and SDE models using MCMC. We place independent uniform priors on each parameter so that 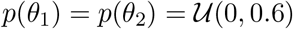 and 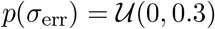. If prior knowledge about the population (i.e., the cell line) is available, perhaps based upon previously conducted experiments, this can be incorporated into the analysis through an informative prior. For example, upper bounds that define reasonable values for biological parameters are routinely applied in this context [90].

In figure 4*a–i*, we show MCMC results for the birth-death process using the ODE model. Based on the structural identifiability results, we expect the likelihood (and for a uniform prior, the posterior density) to be constant along the identifiable parameter combination 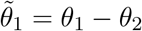, and we see this in figure 4d. These results also suggest that, should one of *θ*_1_ or *θ*_2_ be known (for example, if the cells are treated with an anti-proliferative drug that enforces *θ*_2_ = 0 [144]) the other be identifiable. However, lower and upper bounds for *θ*_1_ and *θ*_2_, respectively, are able to be established as a direct consequence of the prior assumption that all parameters are strictly positive. Examination of univariate credible intervals, shown in table 1, reveals that each parameter cannot individually be identified within 3–4 orders of magnitude, a hallmark of non-identifiability [29]. We note that *σ*_err_ is practically identifiable (figure 4*i*, 95% CrI: (0.1448,0.1907)) from the ODE model, however it will always be overestimated as the observation model for the ODE model must also account for the intrinsic noise of the process.

**Table 1.** 95% credible intervals, and diagnostics, for the parameter estimates for the birth-death process. Credible intervals are approximated using the MCMC quantiles after burn-in.

|  |  | ODE |  |  | SDE |  |  |
| --- | --- | --- | --- | --- | --- | --- | --- |
| | True | 95% CrI | $\hat{R}$ | $S_{\text{eff}}$ | 95% CrI | $\hat{R}$ | $S_{\text{eff}}$ |
| $\theta_1$ | 0.2 | (0.1276,0.5891) | 1.00056 | 2292 | (0.1609,0.4059) | 1.01068 | 104 |
| $\theta_2$ | 0.1 | (0.0130,0.4744) | 1.00056 | 2300 | (0.0477,0.3016) | 1.01107 | 107 |
| $\sigma_{\text{err}}$ | 0.05 | (0.1448,0.1907) | 1.00242 | 2254 | (0.0270,0.0667) | 1.00079 | 364 |

**Figure 4.**
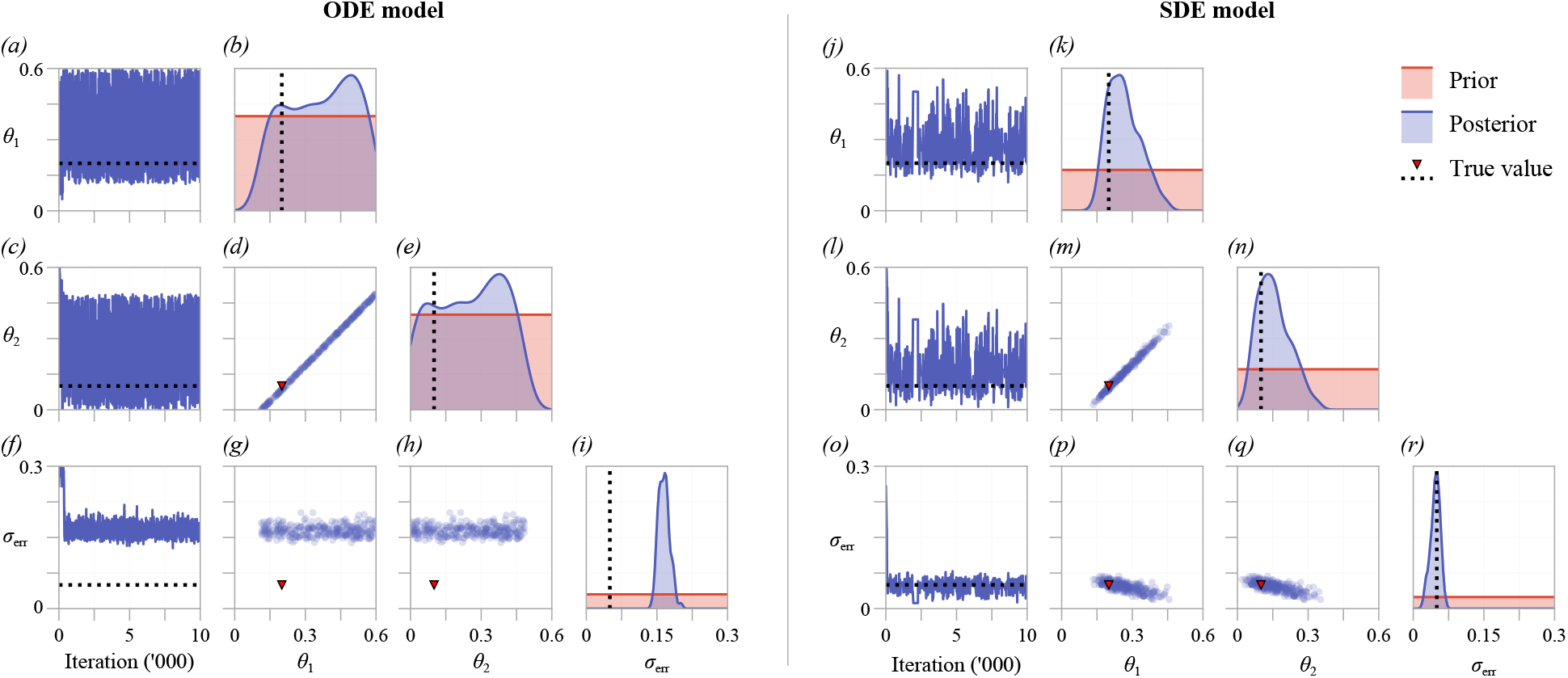
MCMC results for (*a–i*) an ODE and (*j–r*) an SDE description of the birth-death process. (*a,c,f*) and (*j,l,o*) show trace plots for the ODE and SDE models, respectively. Kernel density estimates of the posterior for each parameter ((*b,e,i*) and (*k,n,r*)), and bivariate scatter plots ((*d,g,h*) and (*m,p,q*)), are produced by thinning the MCMC chains by using every 100th sample from four independent MCMC chains, after burn-in.

We repeat the analysis for the SDE model, results of which are shown in figure 4*j–r*. For the prior support chosen, both *θ*_1_ and *θ*_2_ are practically identifiable, as seen in figure 4*k,n*. Further, 95% credible intervals identify each parameter within a single order of magnitude (table 1). While structural identifiability analysis revealed that the SDE model is identifiable in the limit of infinite, noise-free data, it is not necessarily so for data with a realistic signal-to-noise ratio, characterised by the noise model parameter *σ*_err_. In our case, if prior knowledge provided an upper bound for *θ*_1_ and *θ*_2_ at, for example, 0.3, conclusions of practical identifiability may be analogous to those of the ODE model. We see this in table 1, where the upper bounds of the credible intervals for *θ*_1_ and *θ*_2_ extend beyond 0.3. This is also evident from both the bivariate scatter plot (figure 4*m*) and MCMC trace plots (figure 4*j,l*), where posterior samples above 0.3 are regularly drawn for both *θ*_1_ and *θ*_2_. As the SDE explicitly accounts for intrinsic noise, *σ*_err_ is identifiable with estimates close to the true value, in contrast to results from the ODE model.

### 3.2 Two-pool model

Next, we consider partial observations of a process governed by a two-pool model, describing the decay of a substance that is able to transfer between two pools (figure 2*b*). Identifiability of a two-pool model was first examined in the fundamental study of Bellman and Åström [34] as they introduced the concept of structural identifiability. The model can represent, for example, human cholesterol distribution dispersed through two-pools (for example, two organs), where measurements are taken from a tracer in the first pool [104]. Bellman [34] and later Cobelli [35] show that, for an ODE model, the pool transfer and decay rates are not structurally identifiable. We consider practical identifiability for synthetic data comprising noisy measurements of the first pool at 10 equally spaced time points in five identically prepared experiments. Although measurements of the second pool are not taken, we assume, for demonstration purposes, that the initial concentration in each pool is zero before a known amount is introduced to the first pool, thus the full initial condition is known. In practice, the initial condition may also depend on a set of unknown parameters, and we focus on this with the epidemic model.

#### 3.2.1 Model formulation and moment equations

The two-pool model can be expressed as the bio-chemical reaction network

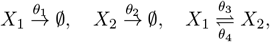

with stoichiometries ***ν***_1_ = (−1, 0)^*T*^, ***ν***_2_ = (0, −1)^*T*^, ***ν***_3_ = (−1,1)^*T*^ and ***ν***_4_ = (1, −1)^*T*^; and propensities *a*_1_(**X**_*t*_) = *θ*_1_*X*_1_, *a*_2_(**X**_*t*_) = *θ*_2_*X*_2_, *a*_3_(**X**_*t*_) = *θ*_3_*X*_1_ and *a*_4_(**X**_*t*_) = *θ*_4_*X*_2_. Here, we denote **X**_*t*_ = (*X*_1,*t*_, *X*_2,*t*_)^*T*^ as the concentration of cholesterol in the first and second pools, respectively. The observed concentration, *Y_t_*, is described by the noise model

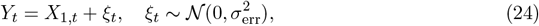

in which we consider that the data are subject to measurement error in the form of additive Gaussian noise [9, 145, 146]. We show 100 realisations of the SSA for the two-pool model in figure 3*b*, and the synthetic data used for practical identifiability analysis in figure 3f. The data are generated using the initial condition **X**_0_ = (100,0)^*T*^ and target parameter values *θ*_1_ = 0.1, *θ*_2_ = 0.2, *θ*_3_ = 0.2, *θ*_4_ = 0.5 and *σ*_err_ = 2. Here, we note that *σ*_err_ ≪ *X_t_* (figure 3*c*), which ensures *Y_t_* > 0.

The CLE for the two-pool model is

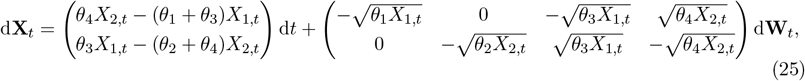

and the moment equations are given to second order by

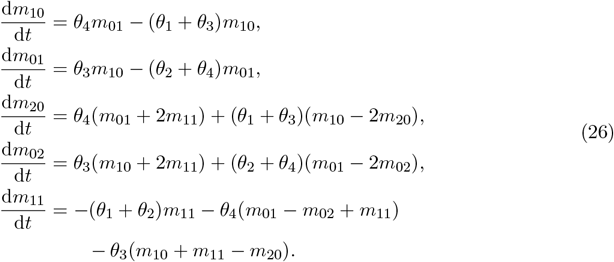

The moments of the observed cholesterol concentration are given in the noise-free limit by *n*_1_ = *m*_10_ and *n*_2_ = *m*_20_. As with the birth-death process, all elements of ***α***(·) and ***σ***(·)***σ***(·)^*T*^ are linear in **X**_*t*_, so the moment equations are closed at every order and, therefore, exact.

#### 3.2.2 Structural identifiability

The two-pool model provides an archetypical example of structural non-identifiability in an ODE model [34,35]. Unless a restriction is placed on one of the parameters (for example, if decay of the substance can only occur from the first pool so *θ*_2_ = 0), the model parameters are structurally non-identifiable: many parameter combinations give identical behaviour in the ODE model. Therefore, the model parameters cannot be uniquely determined from any amount of noise-free experimental data if observations are made from only the first pool.

We assess structural identifiability of an SDE description of the two-pool SDE model using `DAISY` with the system of moment equations up to second order (equation (26)). While the ODE model is structurally non-identifiable, the SDE model is structurally identifiable. Therefore, in the limit of infinite, noise-free data, the model parameters can be uniquely determined from an SDE description of the two-pool model.

#### 3.2.3 Practical identifiability

To assess practical identifiability of the two-pool model, we apply MCMC to infer ***θ*** = (*θ*_1_, *θ*_2_, *θ*_3_, *θ*_4_, *σ*_err_). Initially, independent uniform priors are chosen such that 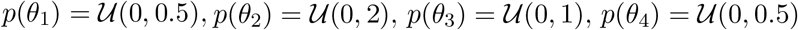, and 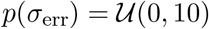. The support of each prior is chosen to cover a range of magnitudes over the target parameter values. Results from four independent pilot chains, each initiated at a random sample from the prior, are shown in figure 5*a–f*. In figure 5*a* we see that the log-likelihood estimate rapidly stabilises, indicating that the chain has moved to a high-likelihood region of the parameter space. Results for *σ*_err_ and *θ*_3_ also rapidly stabilise, indicating that these parameters are practically identifiable [29].

**Figure 5.**
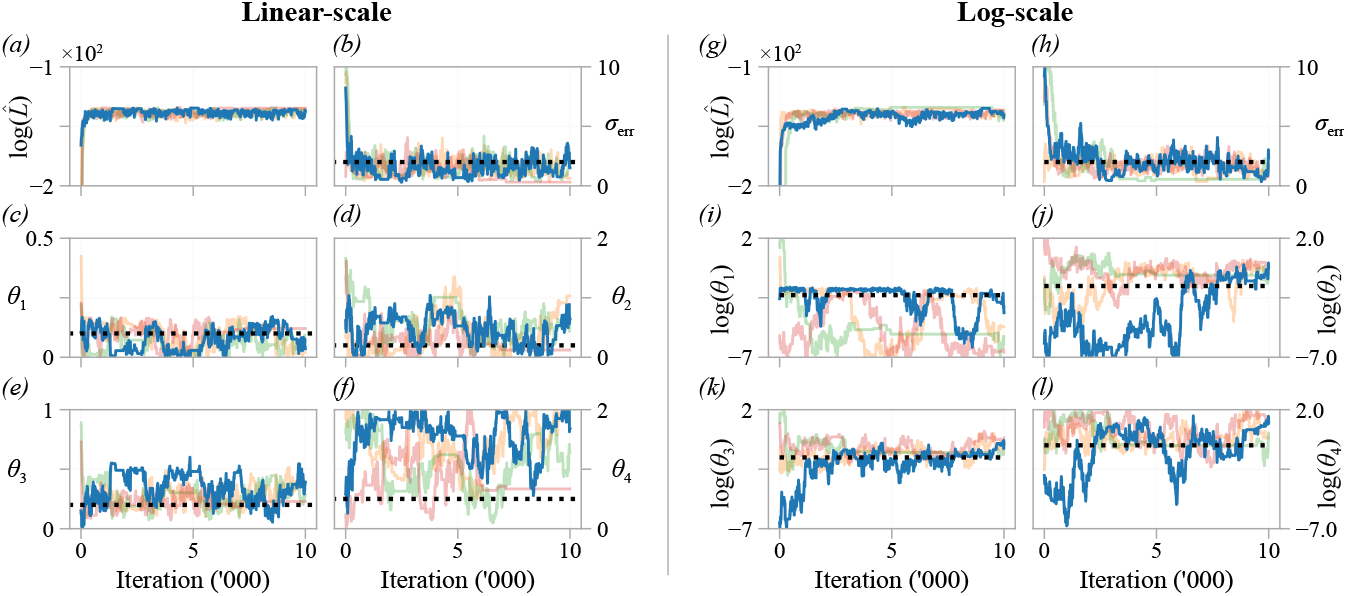
Pilot MCMC trace plots, and log-likelihood estimates, of four chains for the two pool SDE model on with (*a–f*) untransformed parameters; and (*g–l*) transformed parameters. Priors for each parameter are uniform with support corresponding to the respective axis limits. The target parameters, used to generate synthetic data, are indicated (black dashed line).

Results for the remaining three kinetic rate parameters in figure 5*c,d,f* indicate that *θ*_1_, *θ*_2_ and *θ*_4_ are practically non-identifiable. In particular, chains for *θ*_1_ and *θ*_2_ spend a non-negligible time near zero, indicating that the model may be indistinguishable (using the available data) from a model where removal only occurs from a single pool.

We next repeat the analysis using MCMC to infer ***θ***_*_ = (log*θ*_1_, log*θ*_2_, log*θ*_3_, log*θ*_4_, *σ*_err_). Inferring the logarithm of rate parameters will provide more detailed information about the magnitude of rate parameters potentially close to zero [102]. This transformation provides an excellent example of why even a uniform prior is informative, since a uniform prior placed on the linear-scale is not uniform on the log-scale. A uniform prior on the linear-scale makes parameters of a smaller magnitude less likely than a larger magnitude. The priors are again chosen to be independent and uniform (on the log-scale), such that 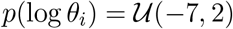 for all *i* and 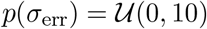 as before. The support of each prior is chosen, again, to cover a range of magnitudes above and below that of the target parameter values. Results in figure 5*k* confirm that *θ*_3_ is practically identifiable, while *θ*_2_ and *θ*_4_ are practically non-identifiable. From results in figure 5*l* we term *θ*_4_ *one-sided identifiable*: the parameter has an identifiable lower bound, and is distinguishable from zero.

To visualise correlations between inferred parameters, we tune the proposal kernel (equation (15)) and run the MCMC algorithm for 30,000 iterations, results are shown in figure 6 and table 2. If only the univariate marginal distributions are considered, all parameters except for *θ*_4_ may be classified as practically identifiable. However, our analysis shows that *θ*_1_ and *θ*_2_ are distinguishable only within a large range of magnitudes. A strong correlation is seen between *θ*_1_ and *θ*_2_, indicating that the total substance exit rate, *θ*_1_ + *θ*_2_, may be practically identifiable. If one of *θ*_1_ or *θ*_2_ were known in advance, perhaps based on past experimental knowledge, the other may become practically identifiable. Further, results from the tuned chains verify that *θ*_3_ is practically identifiable (95% CrI (0.1356,0.4857)) and *θ*_4_ is distinguishable from zero.

**Table 2.** 95% credible intervals, and diagnostics, for the parameter estimates (on the linear-scale) for the two-pool model. Credible intervals are approximated using the MCMC quantiles after burn-in.

| | True | 95% CrI | $\hat{R}$ | $S_{\text{eff}}$ |
| --- | --- | --- | --- | --- |
| $\theta_1$ | 0.1 | (0.0042, 0.1503) | 1.0024 | 510 |
| $\theta_2$ | 0.2 | (0.0307, 1.0699) | 1.0014 | 456 |
| $\theta_3$ | 0.2 | (0.1356, 0.4857) | 1.0023 | 515 |
| $\theta_4$ | 0.5 | (0.4372, 1.9585) | 1.0004 | 741 |
| $\sigma_{\text{err}}$ | 2.0 | (0.5715, 2.8773) | 1.0089 | 409 |

**Figure 6.**
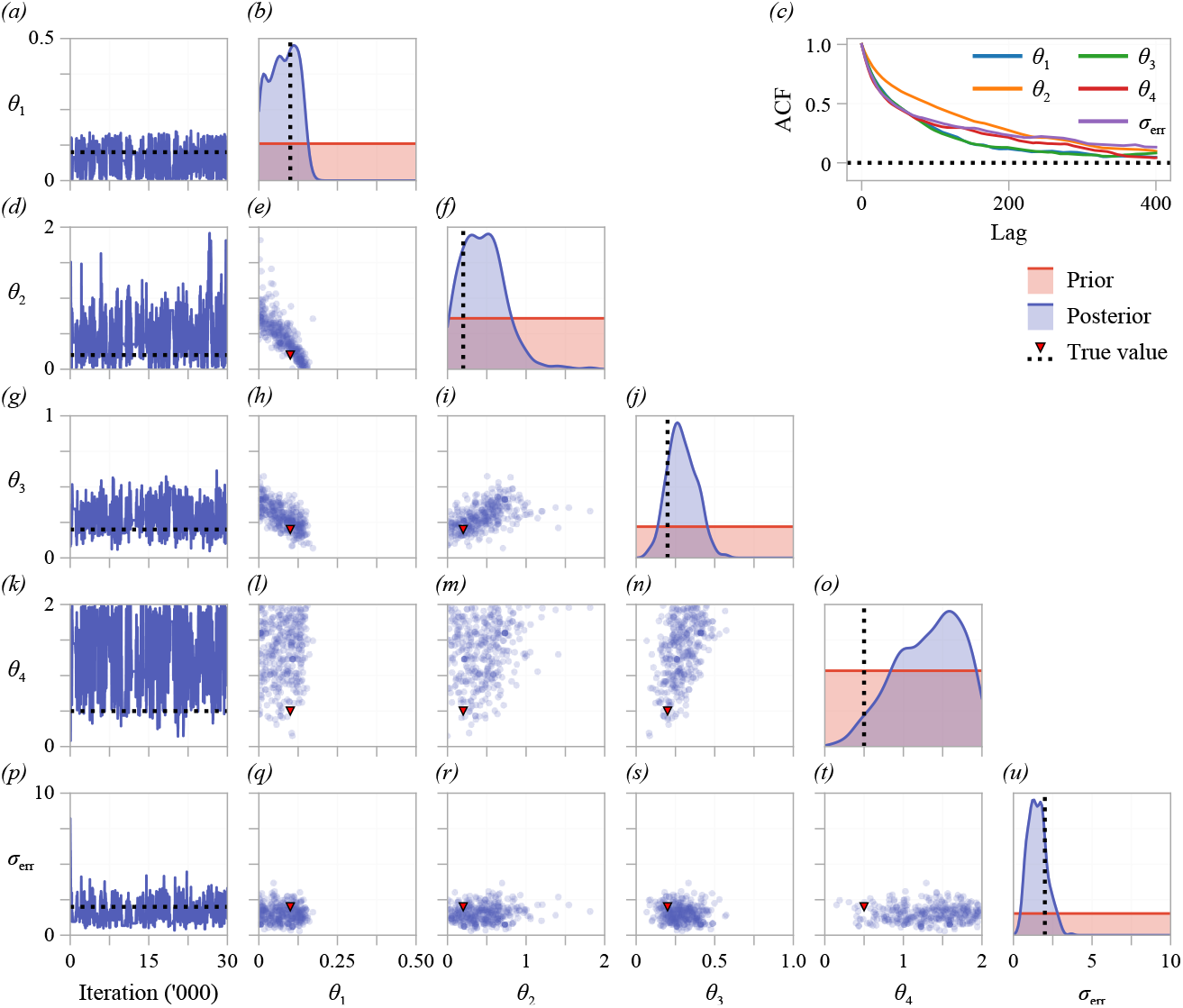
Tuned MCMC results for the two-pool model with a parameters on the linear-scale. The left-most column shows an MCMC trace from a single chain. Kernel density estimates of the marginal posterior for each parameter and bivariate scatter plots are produced using every 300th sample from four independent MCMC chains, after burn-in. The autocorrelation function for a single chain is shown in (*c*), indicating that every 300th sample is approximately independent.

#### 3.3 Epidemic model

Here, we consider a four-compartment epidemic model — the SEIR model [105–107] (figure 2*c*). In this model, susceptible individuals, *S*, are infected due to interactions with infectious individuals, *I*, and undergo an unknown period of time during which they have been exposed, *E*, but are not themselves infectious. Infectious individuals either recover or are removed from the total population, *R*. A noisy unknown proportion, *ξ*, with mean *μ*_obs_, of the number of infectious and recovered individuals is monitored. This captures a testing regime where not all infectious or recovered individuals are tested. We supplement these results by considering a scenario where the same unknown proportion of the exposed individuals is also monitored during the early part of the epidemic.

The kind of data available for the epidemic model differs significantly from that for the experiment-based models we have considered thus far: we are interested in a practical identifiability problem where data from only a single time-series is available, which mirrors data available from an actual epidemic [147]. We first consider practical identifiability using data from the early part of the epidemic, before the number of cases is observed to decrease. Next, these results are compared to a case where data further through the course of the epidemic is considered (figure 3*g*). Initially, 10 infected individuals and 10 recovered individuals are detected. For simplicity we assume there is no noise in these initial observations, so the number of infected and recovered individuals is given by 10/*μ*_obs_. An unknown number of individuals, *E*_0_, are initially exposed. In our analysis, we assume that *E*_0_ is not of direct interest, and we class it a nuisance parameter.

##### 3.3.1 Model formulation and moment equations

The SEIR model can be represented by the following bio-chemical reactions

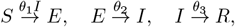

with stoichiometries ***ν***_1_ = (−1,1, 0, 0)^*T*^, ***ν***_2_ = (0, −1, 1, 0)^*T*^ and ***ν***_3_ = (0, 0, −1, 1)^*T*^; and propensities *a*_1_(**X**_*t*_) = *θ*_1_*S_t_I_t_*, *a*_2_(**X**_*t*_) = *θ*_2_*E_t_* and *α*_3_(**X**_*t*_) = *θ*_3_*I_t_*. Here, we denote **X_*t*_** = (*S_t_, E_t_, I_t_, R_t_*)^*T*^ as the number of individuals in each compartment. Two observations are made,

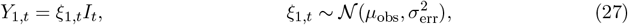

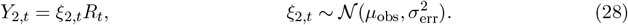

Here, *Y*_1,*t*_ and *Y*_2,*t*_ describe the observed number of infected individuals and recovered individuals, respectively. We further assume that *μ*_obs_, the average observed proportion; and *σ*_err_, the observation error, are unknown and must be estimated. We show 100 realisations of the SSA for the epidemic model in figure 3*c*, and synthetic data used for practical identifiability analysis in figure 3*g*. The data are generated using the initial condition **X**_0_ = (500 – *E*_0_, *E*_0_, 10/*μ*_obs_, 10/*μ*_obs_)^*T*^ and target parameter values *θ*_1_ = 0.01, *θ*_2_ = 0.2, *θ*_3_ = 0.1, *E*_0_ = 20, *μ*_obs_ = 0.5 and *σ*_err_ = 0.05. Here, we note that *σ*_err_ ≪ *μ*_obs_, ensuring that *Y*_1,*t*_ and *Y*_2,*t*_ remain positive.

The moment equations differ from the previous two models considered in that they are not closed. Therefore, the first order moment equations are not equivalent to those for the corresponding ODE model [30], unless a mean-field closure is drawn at first order. To make progress, we close the moment equations after second order to form an approximate system of moment equations for the first two moments. We give the system of 14 moment equations, under all three moment closures considered, as supporting material. The moments of the observation variables are given in the noise-free limit by

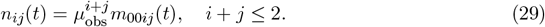

In the supporting material we produce numerical solutions to the moment equations for the epidemic model for each closure considered (figure S1). All closures predict visually identical behaviour at first order, and the pair-approximation and Gaussian closures are in agreement at second order. For the target parameters we consider, the mean-field closure does not agree at second order with the more advanced closures. Whereas a numerical solution to the moment equations for the pair-approximation and Gaussian closures is readily obtainable from a standard solver in `Julia` [148], the mean-field closure required a positivity-preserving Patankar-type method [149] to avoid blow up.

##### 3.3.2 Structural identifiability

We assess structural identifiability of the approximate system of moment equations in `DAISY` and `GenSSI2`, results are shown in table 3. The ODE model, equivalent to a mean-field closure (equation (10)) drawn after the first moment, is structurally non-identifiable. The second-order systems, for all closures, are structurally identifiable (table 3). As the second-order systems are approximate, this analysis is not conclusive for the SDE. However, we can conclude that if the mean and variance of the epidemic model (the first two moments) are modelled using the system of moment equations, and data is available accordingly, the parameters are able to be accurately estimated in the limit of infinite, noise-free data. We highlight the computational cost in `DAISY` of introducing complexity into the moment equations through the closure methods. The pair-wise closure, equation (11), which introduces a quotient, and the Gaussian closure, equation (12), which introduces a cubic, take significantly longer using `DAISY` to assess than the mean-field closure, equation (10), yet give the same result. However, unlike MCMC, we note that structural identifiability results are deterministic, and independent of user choices such as prior, number of particles, and generated or real synthetic data.

**Table 3.** Structural identifiability of the partially observed SEIR model assessed in `DAISY` and `GenSSI2`. Structural identifiability of the SDE is assessed using each closure method for third and higher order moments. Note that the ODE model is equivalent to the SDE model with a mean-field closure for second and higher order moments. Runtimes correspond to a 3.7GHz quad-core i7 desktop machine running Windows 10.

| Model | Structural identifiability | Runtime (DAISY) | Runtime (GenSSI2) |
| --- | --- | --- | --- |
| ODE | Non-Identifiable | 5 seconds | 5 seconds |
| SDE (mean-field closure) | Identifiable | 5 minutes | 2 seconds |
| SDE (pair-wise closure) | Identifiable | 16 hours | 2 seconds |
| SDE (Gaussian closure) | Identifiable | 7 hours | 2 seconds |

##### 3.3.3 Practical identifiability

We assess practical identifiability of the epidemic model using MCMC to infer ***θ*** = (*θ*_1_, *θ*_2_, *θ*_3_, *E*_0_, *μ*_obs_, *σ*_err_). Independent uniform priors are placed on each parameter so that 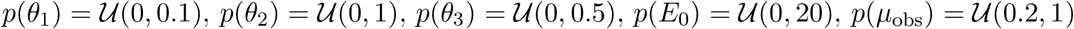 and 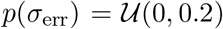. Results are shown in figure 7, where we initiate each chain at the same location for all forms of data we consider.

**Figure 7.**
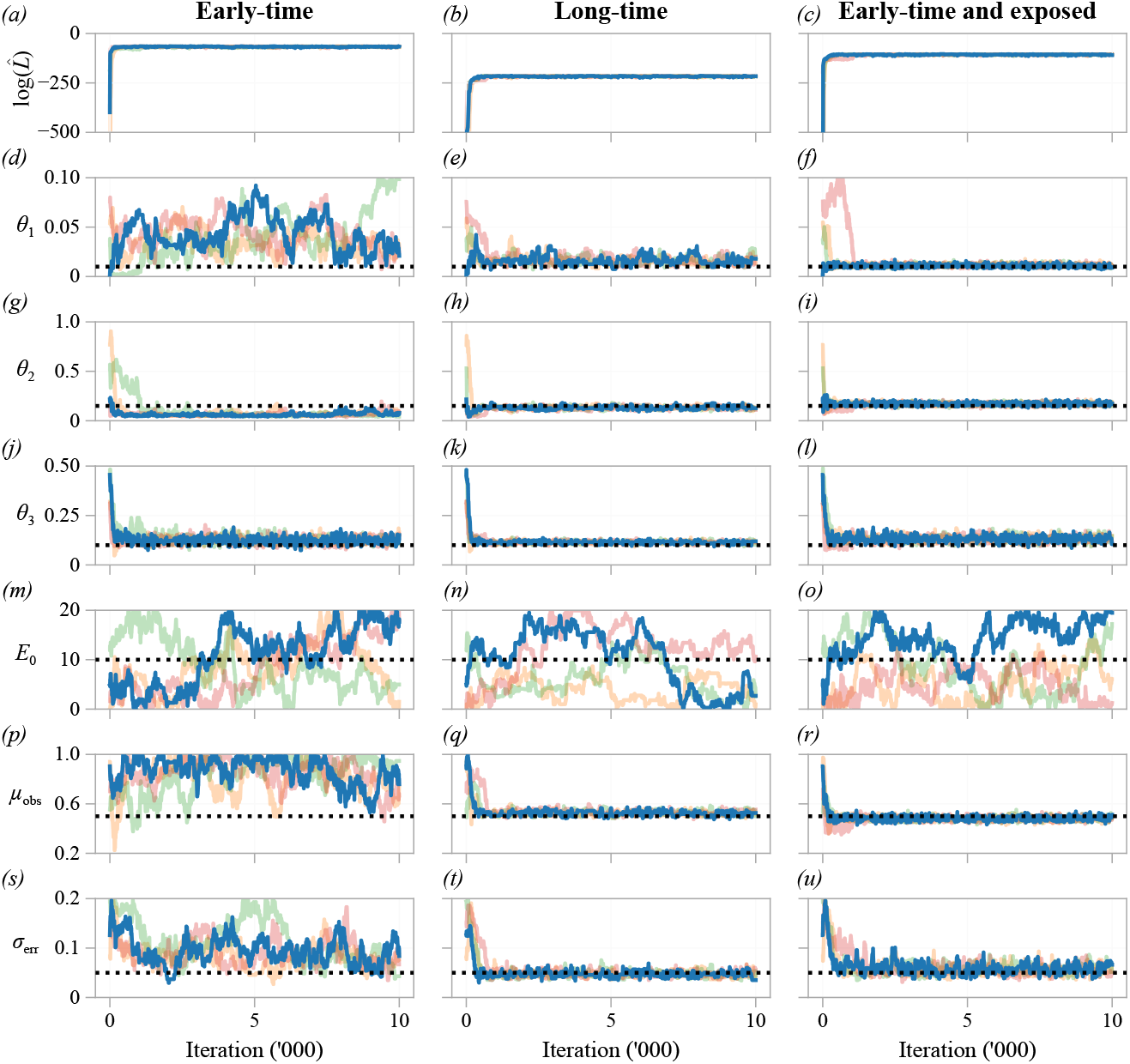
Pilot MCMC trace plots, and log-likelihood estimate, of four chains for the epidemic model. We consider data comprising noisy observations of an unknown proportion of the number of infected and recovered individuals during the early part of the epidemic (first column) and throughout the epidemic (second column). We supplement these results by considering the case we are also able to observe the same unknown proportion of the number of exposed individuals during the early part of the epidemic (third column). Priors for each parameter are uniform with support corresponding to the respective axis limits. The target parameter set, used to generate synthetic data, are indicated (black dashed line).

First, we assess identifiability when only early-time data is available. The log-likelihood estimate rapidly stabilises (figure 7*a*), indicating that the chains have moved to a high-likelihood region of the parameter space [29]. Results for *θ*_3_, the recovery rate, also stabilise, indicating that *θ*_3_ is structurally identifiable. Eventually, we see the estimate for *θ*_2_ stabilises in all chains, however they under-estimate the target value, although proposals equal to and greater than the target value *θ*_2_ = 0.2 are occasionally accepted. To compensate, the estimate of *θ*_1_ stabilises, and covers a region an order of magnitude greater than the target (θ_1_ = 0.01). Therefore, although *θ*_1_ is practically identifiable to a large, but finite, range of values, we classify *θ*_1_ as non-identifiable from the short-time data. Estimates for *E*_0_ and *μ*_obs_ in figure 7*m,p* do not stabilise, and are practically non-identifiable.

Next, we consider a scenario where long-time data are available, such that the number of infected individuals is observed to eventually decrease. The log-likelihood estimate (figure 7*b*) and chains for all parameters, except *E*_0_, are observed to stabilise, indicating that all parameters of interest are now practically identifiable. We supplement these results by considering a third scenario, where only early-time data are available, but the same unknown proportion of the number of exposed individuals is also monitored. As with the long-time data, all parameters of interest are now practically identifiable.

We perform a posterior predictive check [135] of the epidemic model to compare the model prediction—which accounts for parameter uncertainty, intrinsic noise and observation error—to the synthetic data used for inference. We discard the first 3,000 samples from each pilot chain as burn-in, and resample 10,000 parameter combinations for each data type considered. Results in figure 8 show that, in all cases, the model predictions are in agreement with the full time-course (although, we note, the long-time data is only used to calibrate parameters in figure 8*b*). Results in figure 8*a*, for the short-time data, highlight how practical non-identifiability affects model predictions. These results predict an epidemic size at *t* = 30 is noticeably wider and higher than those for the data types where *θ*_1_ is practically identifiable. Further, the lower 95% credible interval for the observed number of infected individuals reduces much faster than that predicted by the other data types.

**Figure 8.**
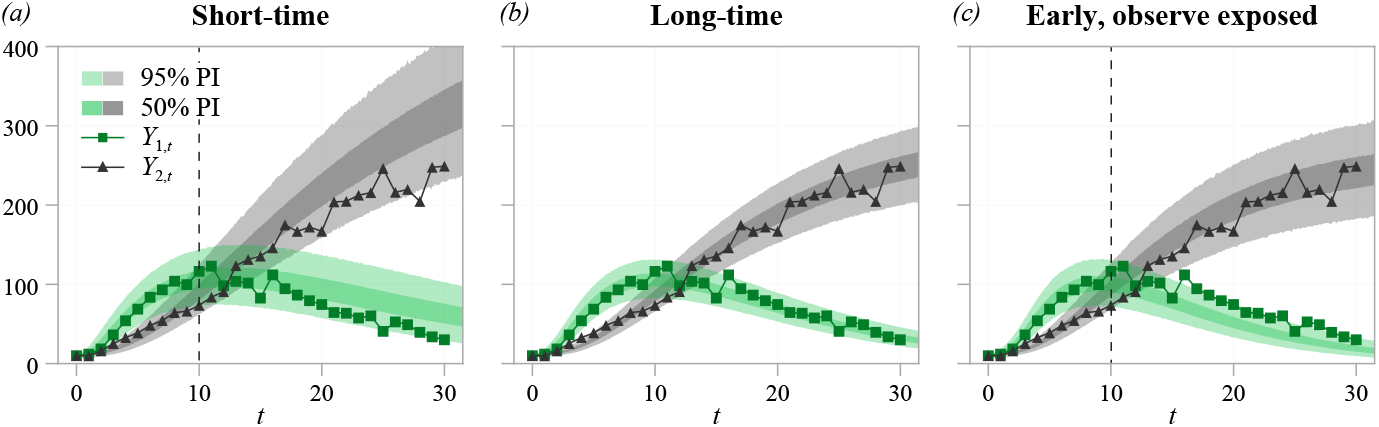
Posterior predictive distribution for the epidemic model using (*a*) short–time data; (*b*) long-time data; and (*c*) short-time data where observations are also made of the number of exposed individuals. In (*a,c*), the dashed line indicates the last observation point used for inference. The first 3,000 samples from each pilot chain is discarded as burn-in. We resample 10,000 parameter combinations (with replacement) and solve the SDE model to estimate posterior predictive intervals (PIs). Shown are 50% (darker) and 95% (lighted) prediction intervals computed from the quantiles of the posterior predictive distribution.

#### 3.4 *β*-insulin-glucose circuit

Finally, we consider a non-linear model of glucose homeostasis, the *β*-insulin-glucose circuit [108,109] (figure 2*d*). Parameterising mathematical models of glucose homeostasis is important for the development of patient-specific insulin delivery for type 1 diabetics [49]. Time-series data of blood glucose concentration is available from continuous glucose monitoring sensors, a critical component of type 1 diabetes management [50,70], an example of which is shown in figure 1*f*. The model describes the regulation of blood plasma glucose by insulin secreted by pancreatic *β* cells. Glucose is introduced into the system through a base production plus a meal intake, *u*(*t*), and decays linearly according to the insulin concentration. Insulin is secreted by *β* cells at a rate given by a non-linear Hill function [109]. *β* cells are produced and decay in a non-response to the glucose concentration. We consider identifiability for synthetic data comprising noisy measurements of the *β* cell and glucose concentrations, but not the insulin concentration. The data consists of five independent experiments, each comprising 15 time-series observations following a meal intake. We only consider inference for two biophysical parameters: *θ*_1_, the insulin secretion rate; and *θ*_2_, the insulin sensitivity. The non-linearities in the model mean that the moment equation approach is not available, and inference using MCMC is computationally expensive. We demonstrate how structural identifiability analysis of the corresponding ODE system [150] can guide analysis of the SDE system and alleviate some of the computational challenges.

##### 3.4.1 Model formulation

We consider a stochastic analogue of the model presented by Karin *et al*. [109]. Denoting **X**_*t*_ = (*β_t_, I_t_, G_t_*)^*T*^ as the concentrations of *β* cells, insulin and glucose, respectively, the propensity functions and corresponding stoichiometries are given by

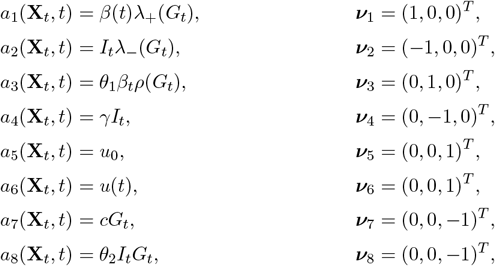

where

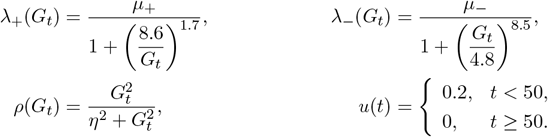

Since *β_t_, I_t_* and *G_t_* denote the concentrations of each substance, and not the population counts, we scale the diffusion term in the CLE to represent the relative concentrations of each substance [57]. Denoting *N_β_, N_I_* and *N_G_* the relative concentration of *β* cells, insulin and glucose, respectively, we write

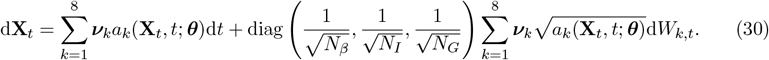

Two observations are made,

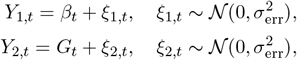

such that *Y*_1,*t*_ and *Y*_2,*t*_ are the observed *β* cell and glucose concentrations, respectively. We show 100 realisations of the SSA for the *β*-insulin-glucose circuit in figure 3*d*, and the synthetic data used for practical identifiability analysis in figure 3*h*. The data are generated using the initial condition **X**_0_ = (322,10, 5)^*T*^ with fixed parameters, *μ*_+_ = 0.21/(24 × 60), *μ*_−_ = 0.025/(24 × 60), *η* = 7.85, *γ* = 0.3, *u*_0_ = 1/30, *c* = 10^−3^, *N_β_* = 1, *N_I_ = N_G_* = 20, and target parameters *θ*_1_ = 0.02, *θ*_2_ = 0.0005 and *σ*_err_ = 0.5 [109]. Here, we note *σ*_err_ ≪ *β_t_,G_t_* (figure 3*d*), which ensures that *Y*_1;*t*_ and *Y*_2,*t*_ remain positive.

##### 3.4.2 Parameter transform

Villaverde *et al*. [151] study structural identifiability of the corresponding ODE model using differential geometry. In the ODE model, *θ*_1_ and *θ*_2_ are structurally non-identifiable, unless the insulin concentration is also observed or one of these two parameters is known. We demonstrate this using MCMC in figure 9*a*, where the marginal posterior for (*θ*_1_, *θ*_2_) covers a hyperbolic region of the parameter space of equal posterior density. In the ODE model, the product *θ*_1_*θ*_2_ is structurally identifiable. To demonstrate this, we perform MCMC on the ODE model with transformed variables 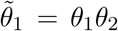 and 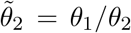, results shown in figure 9*b*. These results also show how inefficient a naïve MCMC proposal can be when correlations between posterior parameters are non-linear. Structural identifiability analysis [151] indicates that the hyperbolic region defined by 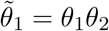 (for a fixed 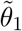) produces indistinguishable behaviour, corresponding to a flat posterior when a uniform prior is applied. Despite this, the tail regions in figure 9*a* are rarely sampled, which could give the impression that the parameters are practically identifiable.

**Figure 9.**
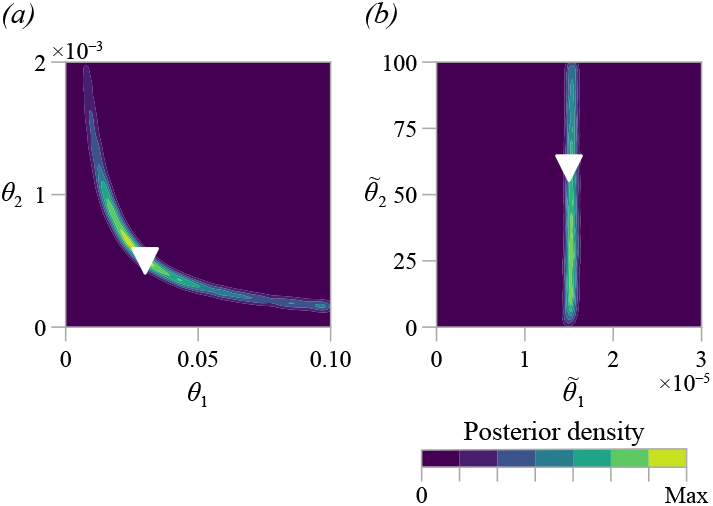
Kernel density of the bivariate marginal posterior distribution of the biophysical parameters in the *β*-insulin-glucose circuit, using the ODE and 100,000 pilot MCMC iterations (the first 3,000 are discarded as burn-in. (a) The posterior for the untransformed parameters, (*θ*_1_, *θ*_2_) shows non-identifiability. (b) The posterior for the transformed parameters, 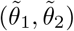, demonstrates that 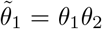 is identifiable, but 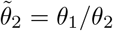 is not.

As the propensity functions for the *β*-insulin-glucose circuit model contain non-polynomial functions, we cannot produce an exact expression for the moment equations. Therefore, we only study practical identifiability using MCMC, and do not consider structural identifiability of the SDE for the *β*-insulin-glucose circuit using the moment equations. Motivated by the structural identifiability analysis of the ODE model, we use MCMC to infer 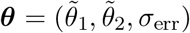, where we only consider the transformed variables 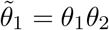 and 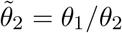.

##### 3.4.3 Practical identifiability

We show MCMC results from four pilot chains in figure 10. The log-likelihood estimate rapidly stabilises (figure 10*a*), as do results for 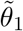 and *σ*_err_ (figure 10*b,d*). As with the ODE model, 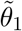 is practically identifiable, but 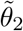 is not. To visualise possible correlations between inferred parameters, we tune the proposal kernel (equation (15)) and run the MCMC algorithm for 10,000 iterations. The univariate marginal distributions, and MCMC trace plots, show that 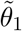 (95% CrI: (1.34,1.67) × 10^−5^) and *σ*_err_ (95% CrI: (0.812,1.049)) are practically identifiable, whereas 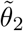 is not (95% CrI: (8.21,97.79)). No large correlations are seen between the parameters 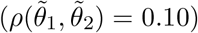, and *θ*_2_ is clearly practically non-identifiable as samples cover the entire range of the prior.

**Figure 10.**
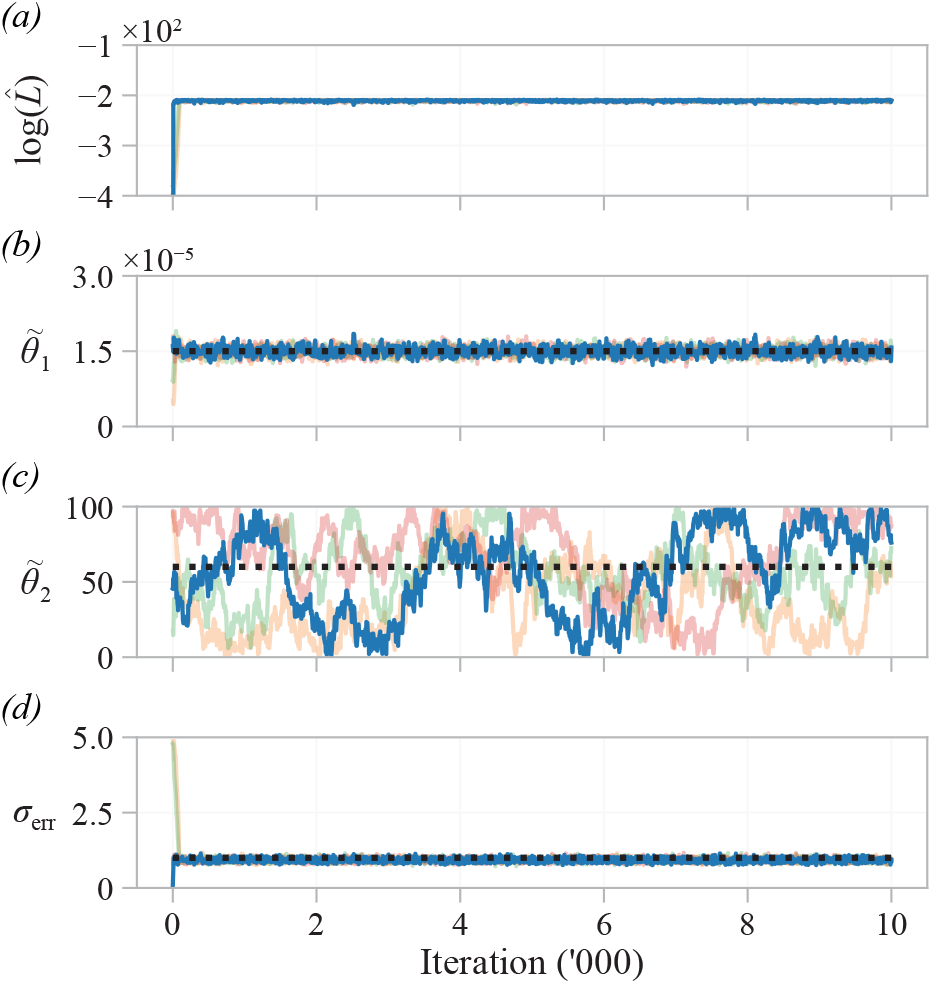
Pilot MCMC trace plots, and log likelihood estimate, of four chains for the *β*-insulin-glucose circuit in the transformed parameter space. The likelihood quickly stabilises, but estimates for 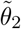 do not, indicating practical non-identifiability. Priors for each parameter are uniform with support corresponding to the respective axis limits. The target parameter set, used to generate synthetic data, are indicated (black dashed line).

**Figure 11.**
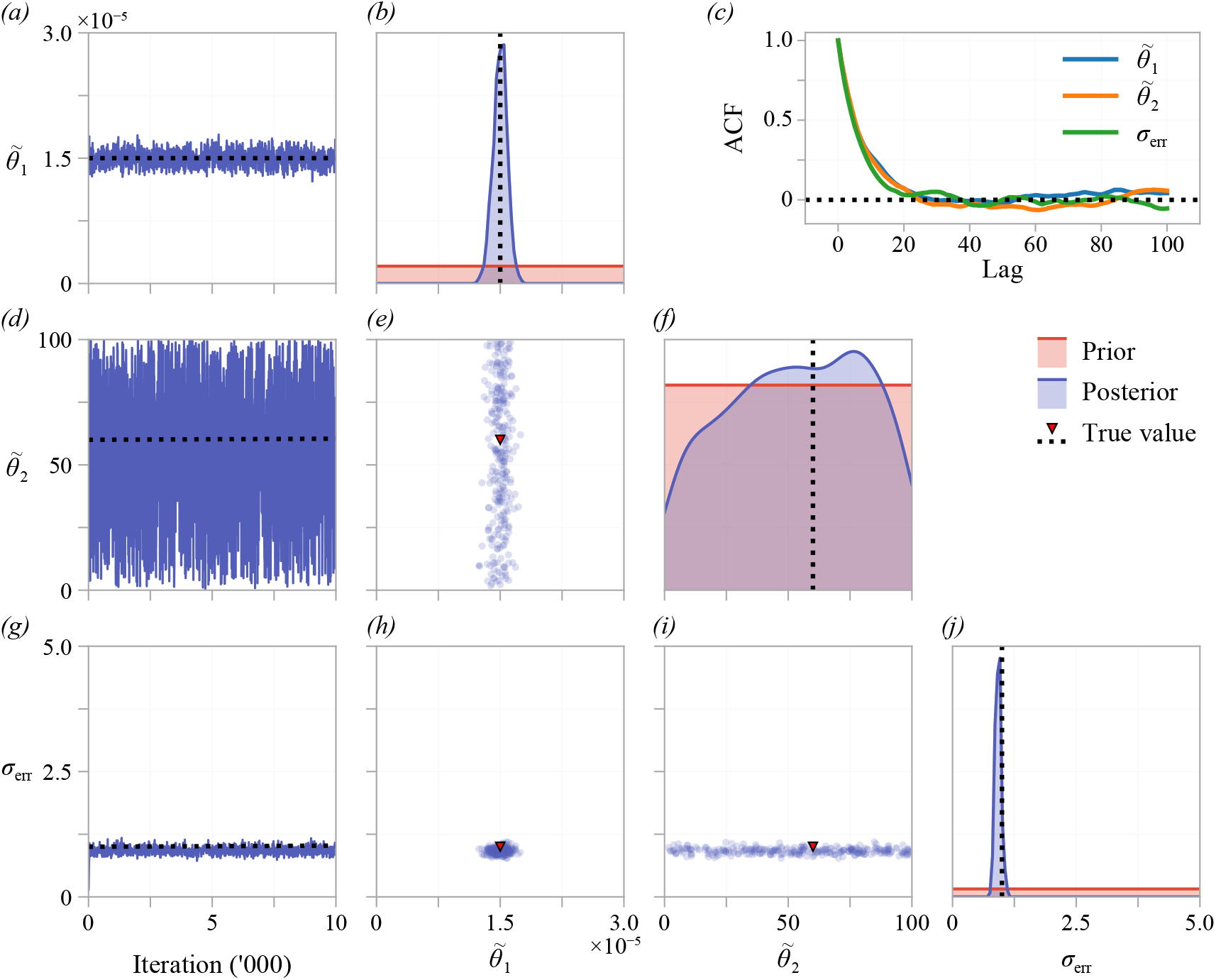
Tuned MCMC results for the *β*-insulin-glucose circuit in the transformed parameter space. The left-most column shows an MCMC trace from a single chain. Kernel density estimates of the marginal posterior for each parameter and bivariate scatter plots are produced using every 100th sample from four independent MCMC chains, after burn-in. The autocorrelation function for a single chain is shown in (*c*), indicating that every 20th sample is approximately independent.

## 4 Discussion

Mathematical models are routinely calibrated to experimental data, with goals ranging from building a predictive model to quantifying biophysical parameters that cannot be directly measured. Much of the usefulness of calibrated models hinges on an assumption that model parameters are identifiable. Heavily over-parameterised models, with large numbers of practically non-identifiable parameters, are often referred to as *sloppy* in the systems biology literature [72,152–154]. Worryingly, these issues of parameter identifiability can often go undetected: models with non-identifiable parameters can still match experimental data (figure 8), but may have poor predictive power and provide little or no mechanistic insight [28]. Identifiability analysis is well-developed for deterministic ODE models, but there is little guidance in the literature to conducting such analysis for the stochastic models that are often vital for interpreting complex experimental data. In this review, we demonstrate how existing techniques can be applied to assess both structural and practical identifiability of SDE models in biology.

### 4.1 Moment dynamics approach

We demonstrate how existing ODE identifiability techniques can be applied directly to stochastic problems by formulating a system of moment equations. In the birth-death process and the two-pool model, the derived moment equations are closed and, therefore, exactly describe the time-evolution of the moments of the SDE. In these two case studies we find that the moment equations are structurally identifiable. This implies structural identifiability of the corresponding SDE model, and parameters can be uniquely estimated in the limit of infinite, noise-free data. For an SDE model this implies an infinite number of *observation-noise* free trajectories of the SDE, since the variability, which relates to higher-order moments, contains information. While we find that the two-pool model of cholesterol distribution is not practically identifiable, establishing structural identifiability is useful as it suggests to the practitioner that the observation process (i.e. observe cholesterol in the first pool) is sufficient, in principle, to fully parameterise the model.

For the epidemic model, the moment equations are not closed, so we study structural identifiability through an approximate system of second-order moment equations. The idea of studying identifiability through an approximate system was first suggested by Pohjanpalo [75], who studies identifiability of ODE systems through a power series expansion. The closed system of moment equations suggest the epidemic SDE model could be structurally identifiable, and these results agree, in our case, with practical identifiability detected using MCMC. More research is needed to establish how identifiability is affected when closing, or truncating, a system of moment equations. For example, if information required to identify model parameters is contained in third or higher order moments, results suggesting that a model is practically non-identifiable from a second order closure will not be indicative of non-identifiability in the SDE model. Furthermore, if structural identifiability differs between moment closures, such a preliminary screening tool needs to be interpreted with caution. If this were the case, a conclusion of structural identifiability is indicative of the model under a particular closure. Recent work suggests that a finite number of moments often contain the information required to identify parameters [155], even for a bimodal distribution and if a closure is applied [156].

Due to the computational constraints placed on analysing structural identifiability of nonpolynomial ODE models, we do not attempt to apply the moment dynamics approach to the stochastic *β*-insulin-glucose circuit model. However, for many models, a mean-field closure corresponds to an ODE description of the system, and studying identifiability of this ODE model can aid practical identifiability analysis of the corresponding SDE. In our case, the corresponding ODE model is structurally non-identifiable due to a hyperbolic relationship between the two parameters of interest: for a fixed *θ*_1_ *θ*_2_, model outputs are indistinguishable [151]. The question of whether an SDE description can provide enough information to practically identify *θ*_1_ and *θ*_2_ can be answered through MCMC, however simple variants of MCMC can struggle when correlations between parameters are strong and non-linear. Therefore, we work in a transformed parameter space where, for the ODE model, 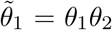 is identifiable but 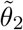 is not (figure 9). This analysis provides a better sense of whether the SDE model captures enough information to identify the parameters, and provides more robust results that are less dependent upon choices made in the MCMC algorithm.

### 4.2 Particle Markov-chain Monte Carlo

We demonstrate practical identifiability by calibrating each model to synthetic data using particle MCMC. We observe the MCMC chains to stabilise in a region of high posterior density, after which time transitions produce samples from the posterior distribution [29]. By visualising MCMC trace plots, we see that estimates of practically identifiable parameters also stabilise, but those of practically non-identifiable parameters do not. These results also demonstrate that, although estimates made of practically identifiable parameters are precise (that is, within a reasonable level of confidence), they are not necessarily accurate. For example, in figure 7*g*, the rate at which exposed individuals become infectious is practically identifiable, but it is underestimated compared to the target value, which could hint at model misspecification.

Given that particle MCMC is computationally expensive, our implementation of a standard technique to detect identifiability from pilot chains carries several advantages. First, pilot chains are regularly generated in the early stages of many inference procedures to establish efficient proposal kernels. Practical identifiability can, therefore, be established as part of an existing workflow. Second, more sophisticated methods are by their very nature more difficult to implement and dependent on practitioner choices, which could obscure results and require more algorithmic experimentation. In comparison, we take an automated approach: aside from the model and choice of prior, the procedure to perform MCMC for each model we consider is identical. Once identifiability is established, the computational cost of MCMC can be alleviated to some extent by adopting a more efficient inference technique. For example, adaptive MCMC [157], sequential Monte-Carlo [158], multi-level methods [159–161], sub-sampling techniques [162] and model-based proposal methods [163] provide significant performance improvements over the standard technique we employ. Further, we expect applying higher-order SDE simulation algorithms, such as a Runge-Kutta method [164], or considering GPU approaches to particle MCMC [165], to improve performance.

As we calibrate to synthetic data for the purpose of a didactic demonstration, we take a pragmatic approach by treating the true values as unknown. Hence, we initiate each chain as a random sample from the prior distribution. This involves a burn-in phase before the MCMC chain settles in an area of high posterior density. For computationally expensive models, such as those found in the cardiac modelling literature [166], synthetic data can be used with pilot chains initiated at the target values. If models have have already been calibrated to experimental data using, for example, maximum likelihood estimation, the chain can be initiated at the calibrated values. MCMC then, relatively cheaply, provides information about the posterior distribution about this point, akin to the Fisher information for models where it can be calculated [41].

MCMC can be applied to detect identifiability for any stochastic model provided an approximation to the likelihood is available. Recent developments to particle MCMC have seen its adoption for more complicated SDE models, such as SDE mixed effects models [167]. For systems with relatively small populations, it may be more appropriate to work directly with an SSA with, for example, a tau-leap method [54, 119]. Alternative approximations to the likelihood, such as those employed by ABC, may be necessary if model complexity requires; for example, should the model include spatial effects [18]. A major drawback of ABC in the context of identifiability is that one must typically decide *a priori* which features of the model and data to match. Common applications of ABC for SDE models match the mean and variance of system [103] or the mean square error between simulations and data [168]. Estimating the likelihood directly, as particle MCMC does, is advantageous when assessing identifiability as it is not clear *a priori* which features of the data and model are significant. For example, some systems might contain the information required for identifiability in higher-order moments or auto-correlations between time-series observations. If ABC is used, a variant that preserves features of the model distribution might be desirable [169].

### 4.3 Modelling noise

In contrast to many studies of identifiability analysis for ODEs, we do not pre-specify parameters in the observation distribution. In a deterministic modelling framework, it is common to assume that all the variability in the data is uncorrelated and sourced from the observation process [41, 90, 170]. Therefore, for an ODE model, the observation parameters can be reliably estimated using the pooled sample variance. For inference on the birth-death ODE model (figure 4), we see that, because the observation variance must now also account for intrinsic noise, the identified value of *σ*_err_ is significantly larger than the target value. For an ODE model with additive homoscedastic Gaussian noise, the posterior mode (in the case of an improper uniform prior), maximum likelihood estimate and least-squares estimate are identical and are independent of the choice of the observed variance. For an SDE model, this is not the case as the intrinsic component of the noise is also modelled implicitly. Therefore, pre-specifying the observation variance could lead to biased estimates and obscure parameter identifiability. We account for this by treating the observation distribution variance as a nuisance parameter that we infer using MCMC, finding it to be identifiable in every case-study considered.

We have focussed our analysis on SDEs derived through the CLE, where the intrinsic noise can provide more information about the process. However, for large populations, the information contained in higher-order moments dissipates: to leading order, 〈*X*^2^〉 → 〈*X*〉^2^ as *X* → ∞. We see this in the epidemic model (figure 3*c*), where the variance is small compared to the scale of the mean. This loss of information in higher-order moments will not be detected by structural identifiability analysis of the moment equations, which is independent of the relative sizes of each moment. As populations become large, the information tends towards that obtained from the equivalent ODE system: this is the assumption behind many mean-field models. There are, however, many other models that contain sources of variability in their own right. For example, Mummert and Otunuga [61] study identifiability of an epidemic model where the infection rate varies according to a white noise process. Other external effects, such as seasonal effects, are often incorporated into epidemic models [171, 172]. In other systems, extrinsic noise describing, for example, the environment, forms a core part of the process and is described by an SDE independent of the population size [48]. Grey-box models use a diffusion term to characterise uncertain physiological effects [63] that could obscure inference, rather than contain information. Making high-level assumptions about which noise process contains information can help with some of the computational challenges by formulating hybrid models containing a mixture of ODEs and SDEs. Particle MCMC carries across, trivially, to any Itô SDE, and the moment equation approach can be applied provided a system of moments be constructed. We have not considered identifiability of SDE models containing non-diffusion noise, such as coloured noise or jump noise. These models lend themselves to different inference techniques, such as forms of rejection sampling [173].

### 4.4 Approaches to computational challenges

The primary computational cost of working with SDE models stems from the need to simulate a suite of trajectories at each iteration of the particle MCMC algorithm. This cost increases not only with the dimensionality of the problem (as for deterministic models) but also with the amount of data, since the number of particles required for an unbiased likelihood estimate increases with the sample size [25,100]. We see this, in particular, when conducting practical identifiability analysis of the two-pool and *β*-insulin-glucose models. These issues have important ramifications for identifiability, as it may not always be feasible to increase the amount of experimental data to rectify practical non-identifiability. Working with a surrogate model, such as a system of moment equations, can help alleviate some of these challenges. For example, establishing structural identifiability—which is requisite for parameter estimation [28]—indicates that the computational investment is worthwhile. Furthermore, such surrogate models can also form a computationally efficient alternative for assessing practical identifiability and performing inference [156,174], while still capturing more information than a purely deterministic description.

A large class of high-dimensional stochastic models lend themselves to structural identifiability analysis through moment equations. For example, CLE descriptions of multi-state ion-channel systems [57] and cascades with many bimolecular reactions [175], can be analysed in terms of a surrogate model using moment equations. This approach can be used because the propensity functions are often polynomial. However, the systems of moment equations are often infinite and require a moment closure approximation to facilitate this analysis. When a moment closure assumption is required, we find that newer software, such as `GenSSI2` in `Matlab`, is computationally advantageous over `DAISY` (table 3). Further, for larger systems, it may not be necessary to consider a full system of second-order moments: a closed system that neglects covariances is also a potentially useful surrogate model.

Many stochastic problems are both computationally expensive to assess using particle MCMC and may not directly permit moment dynamics analysis. The *β*-insulin-glucose model comprises eight reactions and takes approximately 30 hours to perform 10,000 iterations of a pilot chain^2^. These issues are magnified by the quantity of data we consider in this example: five time-series of 15 observations each. Non-polynomial propensity functions mean that an exact expression for the moment equations cannot be derived, so it may not be possible to pre-detect structural non-identifiability through the moment dynamics approach. Fortunately, other approaches, such as those that use polynomial chaos [176], Gaussian processes [177], or the linear noise approximation [93] can provide alternative means of deriving surrogate models. This kind of approximation is already routine in the field of uncertainty quantification, which has deep connections to identifiability [178, 179]. In the future, many of these ideas could allow tractable structural and practical identifiability analysis of large systems of SDEs and, by extension, analysis of spatial problems described by stochastic partial differential equations (SPDEs).

The computational cost of MCMC, in particular for stochastic models with many parameters, has spurred the development of alternatives to explore and exploit the geometry of the likelihood near parameter estimates. Komorowski *et al*. [93] apply the linear noise approximation—which captures first and second order behaviour—to perform sensitivity analysis and calculate the Fisher information matrix for stochastic chemical kinetics models. The concept of *information geometry* [153,180] generalises Fisher information and can be applied to detect identifiability through local information [181], and improve the performance of MCMC algorithms [182]. For SDE models in particular, variational Bayesian techniques provide an efficient alternative to MCMC for parameter estimation [183]. In many cases, mathematical models are calibrated to experimental data to establish the value of a biophysical parameter, not to fully parameterise a model. Profile likelihood [92,184] is widely applied to assess identifiability in ODE models by maximising out parameters that are not of direct interest to reduce the dimensionality of the analysis. Since the bootstrap particle filter that we employ estimates the likelihood function, profile likelihood could be applied to SDE problems.

## 5 Conclusion

It is essential to consider identifiability when performing inference. Yet, there is a scarcity of methods available for assessing identifiability of the stochastic models that are becoming increasingly important. We have provided, through this review, an introduction to identifiability and a guide for performing identifiability analysis of SDE models in systems biology. By formulating a system of moment equations, we show how existing techniques for structural identifiability analysis of ODE models can be applied directly to SDE models [28, 34, 35, 150]. Through synthetic data and particle MCMC, we have demonstrated how to establish practical identifiability for SDE models from data [29, 51].

The analysis we demonstrate is critical for tailoring model complexity to the available data [28]. When a structurally identifiable model is found to be practically non-identifiable, identifiability analysis can guide experiment design to discern the quality and quantity of data required to estimate model parameters [185]. On the other hand, models found to be structurally non-identifiable should be re-parameterised, reduced in complexity, or changed [87,186]. Moving from an ODE to an SDE model can often provide enough information to render an otherwise structurally non-identifiable parameter identifiable: we demonstrate this with the birth-death process. As increasing computing power facilitates inference of complex stochastic models, we expect identifiability to become ever more relevant.

## Data availability

This study contains no experimental data. Code used to produce the numerical results is available as a Julia module on GitHub at github.com/ap-browning/SDE-Identifiability.

## Acknowledgements

This work was supported by the Australian Research Council (DP200100177) and the Air Force Office of Scientific Research (BAA-AFRL-AFOSR-20160007). A.P.B. acknowledges support from the Australian Centre for Excellence in Mathematical and Statistic Frontiers International Mobility Program. R.E.B. would like to thank the Leverhulme Trust for a Leverhulme Research Fellowship, the Royal Society for a Wolfson Research Merit Award, and the BBSRC for funding via BB/R00816/1. The authors thank Oliver Maclaren for helpful discussions, and the three anonymous referees for their comments.

## Author Contributions

All authors designed the research; APB performed the research and wrote the manuscript; APB and DJW implemented the computational algorithms. All authors provided direction, feedback and gave approval for final publication.

## Footnotes

1 Code available on Github at https://github.com/ap-browning/SDE-Identifiability

2 Runtimes for all results produced are available on Github at https://github.com/ap-browning/SDE-Identifiability

## References

[1] Abkowitz JL, Catlin SN, Guttorp P. 1996 Evidence that hematopoiesis may be a stochastic process *in vivo*. Nature Medicine 2, 190–197. (doi:10.1038/nm0296-190).

[2] Elowitz MB, Leibler S. 2000 A synthetic oscillatory network of transcriptional regulators. Nature 403, 335–338. (doi:10.1038/35002125).

[3] Wilkinson DJ. 2009 Stochastic modelling for quantitative description of heterogeneous biological systems. Nature Reviews Genetics 10, 122–133. (doi:10.1038/nrg2509).

[4] Neuert G, Munsky B, Tan RZ, Teytelman L, Khammash M, Oudenaarden Av. 2013 Systematic identification of signal-activated stochastic gene regulation. Science 339, 584–587. (doi:10.1126/science.1231456).

[5] Székely T, Burrage K. 2014 Stochastic simulation in systems biology. Computational and Structural Biotechnology Journal 12, 14–25. (doi:10.1016/j.csbj.2014.10.003).

[6] Kang HW, KhudaBukhsh WR, Koeppl H, Rempała GA. 2019 Quasi-steady-state approximations derived from the stochastic model of enzyme kinetics. Bulletin of Mathematical Biology 81, 1303–1336. (doi:10.1007/s11538-019-00574-4).

[7] Xu B, Kang HW, Jilkine A. 2019 Comparison of deterministic and stochastic regime in a model for Cdc42 oscillations in fission yeast. Bulletin of Mathematical Biology 81, 1268–1302. (doi:10.1007/s11538-019-00573-5).

[8] Swameye I, Muller TG, Timmer J, Sandra O, Klingmuller U. 2003 Identification of nucleocytoplasmic cycling as a remote sensor in cellular signaling by databased modeling. Proceedings of the National Academy of Sciences 100, 1028–1033. (doi:10.1073/pnas.0237333100).

[9] Heron EA, Finkenstädt B, Rand DA. 2007 Bayesian inference for dynamic transcriptional regulation; the Hes1 system as a case study. Bioinformatics 23, 2596–2603. (doi:10.1093/bioinformatics/btm367).

[10] Locke JCW, Elowitz MB. 2009 Using movies to analyse gene circuit dynamics in single cells. Nature Reviews Microbiology 7, 383–392. (doi:10.1038/nrmicro2056).

[11] Young JW, Locke JCW, Altinok A, Rosenfeld N, Bacarian T, Swain PS, Mjolsness E, Elowitz MB. 2011 Measuring single-cell gene expression dynamics in bacteria using fluorescence time-lapse microscopy. Nature Protocols 7, 80–88. (doi:10.1038/nprot.2011.432).

[12] Cho H, Rockne RC. 2019 Mathematical modeling with single-cell sequencing data. bioRxiv p. 710640. (doi:10.1101/710640).

[13] Ritchie K, Shan XY, Kondo J, Iwasawa K, Fujiwara T, Kusumi A. 2005 Detection of non-Brownian diffusion in the cell membrane in single molecule tracking. Biophysical Journal 88, 2266–2277. (doi:10.1529/biophysj.104.054106).

[14] Isaacson SA. 2008 Relationship between the reaction–diffusion master equation and particle tracking models. Journal of Physics A: Mathematical and Theoretical 41, 065003. (doi:10.1088/1751-8113/41/6/065003).

[15] Rienzo CD, Piazza V, Gratton E, Beltram F, Cardarelli F. 2014 Probing short-range protein Brownian motion in the cytoplasm of living cells. Nature Communications 5, 5891. (doi:10.1038/ncomms6891).

[16] Schnoerr D, Grima R, Sanguinetti G. 2016 Cox process representation and inference for stochastic reaction–diffusion processes. Nature Communications 7, 11729. (doi:10.1038/ncomms11729).

[17] Brückner DB, Ronceray P, Broedersz CP. 2020 Inferring the dynamics of underdamped stochastic systems. Physical Review Letters 125, 058103. (doi:10.1103/physrevlett.125.058103).

[18] Browning AP, Jin W, Plank MJ, Simpson MJ. 2020 Identifying density-dependent interactions in collective cell behaviour. Journal of The Royal Society Interface 17, 20200143. (doi:10.1098/rsif.2020.0143).

[19] Golightly A, Wilkinson DJ. 2006 Bayesian sequential inference for stochastic kinetic biochemical network models. Journal of Computational Biology 13, 838–851. (doi:10.1089/cmb.2006.13.838).

[20] Toni T, Welch D, Strelkowa N, Ipsen A, Stumpf MPH. 2009 Approximate Bayesian computation scheme for parameter inference and model selection in dynamical systems. Journal of The Royal Society Interface 6, 187–202. (doi:10.1098/rsif.2008.0172).

[21] Liepe J, Kirk P, Filippi S, Toni T, Barnes CP, Stumpf MPH. 2014 A framework for parameter estimation and model selection from experimental data in systems biology using approximate Bayesian computation. Nature Protocols 9, 439–456. (doi:10.1038/nprot.2014.025).

[22] Schnoerr D, Sanguinetti G, Grima R. 2017 Approximation and inference methods for stochastic biochemical kinetics—a tutorial review. Journal of Physics A: Mathematical and Theoretical 50, 093001. (doi:10.1088/1751-8121/aa54d9).

[23] Bressloff PC. 2017 Stochastic switching in biology: from genotype to phenotype. Journal of Physics A: Mathematical and Theoretical 50, 133001. (doi:10.1088/1751-8121/aa5db4).

[24] Bosco DB, Kenworthy R, Zorio DAR, Sang QXA. 2015 Human mesenchymal stem cells are resistant to paclitaxel by adopting a non-proliferative fibroblastic state. PLOS One 10, e0128511. (doi:10.1371/journal.pone.0128511).

[25] Wilkinson DJ. 2012 Stochastic Modelling for Systems Biology. Boca Raton, Florida: CRC Press 2 edition.

[26] Liao S, Vejchodský T, Erban R. 2015 Tensor methods for parameter estimation and bifurcation analysis of stochastic reaction networks. Journal of The Royal Society Interface 12, 20150233. (doi:10.1098/rsif.2015.0233).

[27] Gutenkunst RN, Waterfall JJ, Casey FP, Brown KS, Myers CR, Sethna JP. 2007 Universally sloppy parameter sensitivities in systems biology models. PLOS Computational Biology 3, e189. (doi:10.1371/journal.pcbi.0030189).

[28] Raue A, Kreutz C, Maiwald T, Bachmann J, Schilling M, Klingmüller U, Timmer J. 2009 Structural and practical identifiability analysis of partially observed dynamical models by exploiting the profile likelihood. Bioinformatics 25, 1923–1929. (doi:10.1093/bioinformatics/btp358).

[29] Hines KE, Middendorf TR, Aldrich RW. 2014 Determination of parameter identifiability in nonlinear biophysical models: A Bayesian approach. The Journal of General Physiology 143, 401–4 16. (doi:10.1085/jgp.201311116).

[30] Roosa K, Chowell G. 2019 Assessing parameter identifiability in compartmental dynamic models using a computational approach: application to infectious disease transmission models. Theoretical Biology and Medical Modelling 16, 1. (doi:10.1186/s12976-018-0097-6).

[31] Kreutz C, Raue A, Timmer J. 2012 Likelihood based observability analysis and confidence intervals for predictions of dynamic models. BMC Systems Biology 6, 120. (doi:10.1186/1752-0509-6-120).

[32] Cedersund G. 2016 Prediction uncertainty estimation despite unidentifiability: an overview of recent developments. In Uncertainty in Biology pp. 449–466. Cham: Springer International Publishing. (doi:10.1007/978-3-319-21296-8_17).

[33] Villaverde AF, Raimúndez E, Hasenauer J, Banga JR. 2019 A comparison of methods for quantifying prediction uncertainty in systems biology. IFAC-PapersOnLine 52, 45–51. (doi:10.1016/j.ifacol.2019.12.234).

[34] Bellman R, Åström K. 1970 On structural identifiability. Mathematical Biosciences 7, 329–339. (doi:10.1016/0025-5564(70)90132-X).

[35] Cobelli C, DiStefano JJ. 1980 Parameter and structural identifiability concepts and ambiguities: a critical review and analysis. American Journal of Physiology-Regulatory, Integrative and Comparative Physiology 239, 7–24. (doi:10.1152/ajpregu.1980.239.1.r7).

[36] Walter E. 1987 Identifiability of Parametric Models. London, United Kingdom: Elsevier Science & Technology. (doi:10.1016/C2013-0-03836-4).

[37] Jaqaman K, Danuser G. 2006 Linking data to models: data regression. Nature Reviews Molecular Cell Biology 7, 813–819. (doi:10.1038/nrm2030).

[38] Bellu G, Saccomani MP, Audoly S, D’Angiò L. 2007 DAISY: A new software tool to test global identifiability of biological and physiological systems. Computer Methods and Programs in Biomedicine 88, 52–61. (doi:10.1016/j.cmpb.2007.07.002).

[39] Miao H, Xia X, Perelson AS, Wu H. 2011 On identifiability of nonlinear ODE models and applications in viral dynamics. SIAM Review 53, 3–39. (doi:10.1137/090757009).

[40] Eisenberg MC, Hayashi MA. 2014 Determining identifiable parameter combinations using subset profiling. Mathematical Biosciences 256, 116–126. (doi:10.1016/j.mbs.2014.08.008).

[41] Daly AC, Gavaghan D, Cooper J, Tavener S. 2018 Inference-based assessment of parameter identifiability in nonlinear biological models. Journal of The Royal Society Interface 15, 20180318. (doi:10.1098/rsif.2018.0318).

[42] Raj A, Oudenaarden Av. 2008 Nature, nurture, or chance: stochastic gene expression and its consequences. Cell 135, 216–226. (doi:10.1016/j.cell.2008.09.050).

[43] Balázsi G, van Oudenaarden A, Collins J. 2011 Cellular decision making and biological noise: from microbes to mammals. Cell 144, 910–925. (doi:10.1016/j.cell.2011.01.030).

[44] Bar-Joseph Z, Gitter A, Simon I. 2012 Studying and modelling dynamic biological processes using time-series gene expression data. Nature Reviews Genetics 13, 552–564. (doi:10.1038/nrg3244).

[45] Ruess J, Lygeros J. 2015 Moment-based methods for parameter inference and experiment design for stochastic biochemical reaction networks. ACM Transactions on Modeling and Computer Simulation 25, 8. (doi:10.1145/2688906).

[46] Soltani M, Vargas-Garcia CA, Antunes D, Singh A. 2016 Intercellular variability in protein levels from stochastic expression and noisy cell cycle processes. PLOS Computational Biology 12, e1004972. (doi:10.1371/journal.pcbi.1004972).

[47] Smith S, Grima R. 2018 Single-cell variability in multicellular organisms. Nature Communications 9, 345. (doi:10.1038/s41467-017-02710-x).

[48] Browning AP, Sharp JA, Mapder T, Baker CM, Burrage K, Simpson MJ. 2020 Persistence as an optimal hedging strategy. bioRxiv. (doi:10.1101/2019.12.19.883645).

[49] Hovorka R, Canonico V, Chassin LJ, Haueter U, Massi-Benedetti M, Federici MO, Pieber TR, Schaller HC, Schaupp L, Vering T, Wilinska ME. 2004 Nonlinear model predictive control of glucose concentration in subjects with type 1 diabetes. Physiological Measurement 25, 905–920. (doi:10.1088/0967-3334/25/4/010).

[50] Facchinetti A. 2016 Continuous glucose monitoring sensors: past, present and future algorithmic challenges. Sensors 16, 1–12. (doi:10.3390/s16122093).

[51] Siekmann I, Sneyd J, Crampin E. 2012 MCMC can detect nonidentifiable models. Biophysical Journal 103, 2275–2286. (doi:10.1016/j.bpj.2012.10.024).

[52] Choi B, Rempała GA, Kim JK. 2017 Beyond the Michaelis-Menten equation: accurate and efficient estimation of enzyme kinetic parameters. Scientific Reports 7, 17018. (doi:10.1038/s41598-017-17072-z).

[53] Turelli M. 1977 Random environments and stochastic calculus. Theoretical Population Biology 12, 140–178. (doi:10.1016/0040-5809(77)90040-5).

[54] Turner TE, Schnell S, Burrage K. 2004 Stochastic approaches for modelling *in vivo* reactions. Computational Biology and Chemistry 28, 165–178. (doi:10.1016/j.compbiolchem.2004.05.001).

[55] Ruess J, Milias-Argeitis A, Lygeros J. 2013 Designing experiments to understand the variability in biochemical reaction networks. Journal of The Royal Society Interface 10, 20130588. (doi:10.1098/rsif.2013.0588).

[56] Parsons TL, Lambert A, Day T, Gandon S. 2018 Pathogen evolution in finite populations: slow and steady spreads the best. Journal of The Royal Society Interface 15, 20180135. (doi:10.1098/rsif.2018.0135).

[57] Dangerfield CE, Kay D, Burrage K. 2012 Modeling ion channel dynamics through reflected stochastic differential equations. Physical Review E 85, 051907. (doi:10.1103/physreve.85.051907).

[58] Dangerfield CE, Kay D, Burrage K. 2011 Comparison of continuous and discrete stochastic ion channel models. 2011 Annual International Conference of the IEEE Engineering in Medicine and Biology Society 2011, 704–707. (doi:10.1109/iembs.2011.6090159).

[59] Gillespie DT. 2000 The chemical Langevin equation. The Journal of Chemical Physics 113, 297–306. (doi:10.1063/1.481811).

[60] Hidalgo J, Pigolotti S, Muñoz MA. 2015 Stochasticity enhances the gaining of bet-hedging strategies in contact-process-like dynamics. Physical Review E 91, 032114. (doi:10.1103/physreve.91.032114).

[61] Mummert A, Otunuga OM. 2019 Parameter identification for a stochastic SEIRS epidemic model: case study influenza. Journal of Mathematical Biology 79, 705–729. (doi:10.1007/s00285-019-01374-z).

[62] Kristensen NR, Madsen H, Jørgensen SB. 2004 Parameter estimation in stochastic grey-box models. Automatica 40, 225–237. (doi:10.1016/j.automatica.2003.10.001).

[63] Duun-Henriksen AK, Schmidt S, Røge RM, Møller JB, Nørgaard K, Jørgensen JB, Madsen H. 2013 Model identification using stochastic differential equation grey-box models in diabetes. Journal of Diabetes Science and Technology 7, 431–440. (doi:10.1177/193229681300700220).

[64] Munsky B, Trinh B, Khammash M. 2009 Listening to the noise: random fluctuations reveal gene network parameters. Molecular Systems Biology 5, 318. (doi:10.1038/msb.2009.75).

[65] Leander J, Lundh T, Jirstrand M. 2014 Stochastic differential equations as a tool to regularize the parameter estimation problem for continuous time dynamical systems given discrete time measurements. Mathematical Biosciences 251, 54–62. (doi:10.1016/j.mbs.2014.03.001).

[66] Enciso G, Kim J. 2019 Embracing noise in chemical reaction networks. Bulletin of Mathematical Biology 81, 1261–1267. (doi:10.1007/s11538-019-00575-3).

[67] Enciso G, Erban R, Kim J. 2020 Identifiability of stochastically modelled reaction networks. arXiv. https://arxiv.org/abs/2006.02272.

[68] Browning AP, McCue SW, Binny RN, Plank MJ, Shah ET, Simpson MJ. 2018 Inferring parameters for a lattice-free model of cell migration and proliferation using experimental data. Journal of Theoretical Biology 437, 251–260. (doi:10.1016/j.jtbi.2017.10.032).

[69] University, Center for Systems Science and Engineering (CSSE) at Johns Hopkins. 2020 COVID-19 data repository. https://github.com/CSSEGISandData/COVID-19. Accessed: 7th July 2020.

[70] Vigers T, Chan CL, Snell-Bergeon J, Bjornstad P, Zeitler PS, Forlenza G, Pyle L. 2019 cgmanalysis: an R package for descriptive analysis of continuous glucose monitor data. PLOS One 14, e0216851. (doi:10.1371/journal.pone.0216851).

[71] Hengl S, Kreutz C, Timmer J, Maiwald T. 2007 Data-based identifiability analysis of non-linear dynamical models. Bioinformatics 23, 2612–2618. (doi:10.1093/bioinformatics/btm382).

[72] Chis OT, Banga JR, Balsa-Canto E. 2011 Structural identifiability of systems biology models: a critical comparison of methods. PLOS One 6, e27755. (doi:10.1371/journal.pone.0027755).

[73] Janzén DLI, Bergenholm L, Jirstrand M, Parkinson J, Yates J, Evans ND, Chappell MJ. 2016 Parameter identifiability of fundamental pharmacodynamic models. Frontiers in Physiology 7, 590. (doi:10.3389/fphys.2016.00590).

[74] Reiersøl O. 1950 Identifiability of a linear relation between variables which are subject to error. Econometrica 18, 375. (doi:10.2307/1907835).

[75] Pohjanpalo H. 1978 System identifiability based on the power series expansion of the solution. Mathematical Biosciences 41, 21–33. (doi:10.1016/0025-5564(78)90063-9).

[76] White LJ, Evans ND, Lam TJGM, Schukken YH, Medley GF, Godfrey KR, Chappell MJ. 2001 The structural identifiability and parameter estimation of a multispecies model for the transmission of mastitis in dairy cows. Mathematical Biosciences 174, 77–90. (doi:10.1016/s0025-5564(01)00080-3).

[77] Maclaren OJ, Nicholson R. 2019 What can be estimated? Identifiability, estimability, causal inference and ill-posed inverse problems. arXiv. https://arxiv.org/abs/1904.02826.

[78] Margaria G, Riccomagno E, Chappell MJ, Wynn HP. 2001 Differential algebra methods for the study of the structural identifiability of rational function state-space models in the biosciences. Mathematical Biosciences 174, 1–26. (doi:10.1016/s0025-5564(01)00079-7).

[79] Saccomani MP, Audoly S, D’Angiò L. 2003 Parameter identifiability of nonlinear systems: the role of initial conditions. Automatica 39, 619–632. (doi:10.1016/S0005-1098(02)00302-3).

[80] Brouwer AF, Eisenberg MC. 2018 The underlying connections between identifiability, active subspaces, and parameter space dimension reduction. arXiv. https://arxiv.org/abs/1802.05641.

[81] Jacquez JA, Greif P. 1985 Numerical parameter identifiability and estimability: Integrating identifiability, estimability, and optimal sampling design. Mathematical Biosciences 77, 201–227. (doi:10.1016/0025-5564(85)90098-7).

[82] Saccomani MP, Thomaseth K. 2018 The union between structural and practical identifiability makes strength in reducing oncological model complexity: a case study. Complexity 2018, 1–10. (doi:10.1155/2018/2380650).

[83] Raue A, Karlsson J, Saccomani MP, Jirstrand M, Timmer J. 2014 Comparison of approaches for parameter identifiability analysis of biological systems. Bioinformatics 30, 1440–1448. (doi:10.1093/bioinformatics/btu006).

[84] Meshkat N, Kuo CEZ, DiStefano J. 2014 On finding and using identifiable parameter combinations in nonlinear dynamic systems biology models and COMBOS: a novel web implementation. PLOS One 9, e110261. (doi:10.1371/journal.pone.0110261).

[85] Villaverde AF, Barreiro A, Papachristodoulou A. 2016 Structural identifiability of dynamic systems biology models. PLOS Computational Biology 12, e1005153. (doi:10.1371/journal.pcbi.1005153).

[86] Hong H, Ovchinnikov A, Pogudin G, Yap C. 2019 SIAN: software for structural identifiability analysis of ODE models. Bioinformatics 35, 2873–2874. (doi:10.1093/bioinformatics/bty1069).

[87] Brouwer AF, Meza R, Eisenberg MC. 2017 Parameter estimation for multistage clonal expansion models from cancer incidence data: A practical identifiability analysis. PLOS Computational Biology 13, e1005431. (doi:10.1371/journal.pcbi.1005431).

[88] Johnston ST, Ross JV, Binder BJ, McElwain DLS, Haridas P, Simpson MJ. 2016 Quantifying the effect of experimental design choices for *in vitro* scratch assays. Journal of Theoretical Biology 400, 19–31. (doi:10.1016/j.jtbi.2016.04.012).

[89] Warne DJ, Baker RE, Simpson MJ. 2017 Optimal quantification of contact inhibition in cell populations. Biophysical Journal 113, 1920–1924. (doi:10.1016/j.bpj.2017.09.016).

[90] Simpson MJ, Baker RE, Vittadello ST, Maclaren OJ. 2020 Practical parameter identifiability for spatio-temporal models of cell invasion. Journal of The Royal Society Interface 17, 20200055. (doi:10.1098/rsif.2020.0055).

[91] Lehmann EL, Fienberg S, Casella G. 1998 Theory of Point Estimation. Secaucus: Springer 2 edition. (doi:10.1007/b98854).

[92] Murphy SA, Vaart AWVD. 2000 On profile likelihood. Journal of the American Statistical Association 95, 449–465. (doi:10.1080/01621459.2000.10474219).

[93] Komorowski M, Costa MJ, Rand DA, Stumpf MPH. 2011 Sensitivity, robustness, and identifiability in stochastic chemical kinetics models. Proceedings of the National Academy of Sciences 108, 8645–8650. (doi:10.1073/pnas.1015814108).

[94] Tavaré S, Balding DJ, Griffiths RC, Donnelly P. 1997 Inferring coalescence times from DNA sequence data. Genetics 145, 505–518. (pmid:9071603).

[95] Pritchard JK, Seielstad MT, Perez-Lezaun A, Feldman MW. 1999 Population growth of human Y chromosomes: a study of Y chromosome microsatellites. Molecular Biology and Evolution 16, 1791–1798. (doi:10.1093/oxfordjournals.molbev.a026091).

[96] Beaumont MA, Zhang W, Balding DJ. 2002 Approximate Bayesian computation in population genetics. Genetics 162, 2025–2035. (pmid:12524368).

[97] Sunnåker M, Busetto AG, Numminen E, Corander J, Foll M, Dessimoz C. 2013 Approximate Bayesian computation. PLOS Computational Biology 9, e1002803. (doi:10.1371/journal.pcbi.1002803).

[98] Wilkinson RD. 2013 Approximate Bayesian computation (ABC) gives exact results under the assumption of model error. Statistical applications in genetics and molecular biology 12, 129–141. (doi:10.1515/sagmb-2013-0010).

[99] Beaumont MA. 2003 Estimation of population growth or decline in genetically monitored populations. Genetics 164, 1139–1160. (pmid:12871921).

[100] Andrieu C, Roberts GO. 2009 The pseudo-marginal approach for efficient Monte Carlo computations. The Annals of Statistics 37, 697–725. (doi:10.1214/07-aos574).

[101] Andrieu C, Doucet A, Holenstein R. 2010 Particle Markov chain Monte Carlo methods. Journal of the Royal Statistical Society: Series B (Statistical Methodology) 72, 269–342. (doi:10.1111/j.1467-9868.2009.00736.x).

[102] Golightly A, Wilkinson DJ. 2011 Bayesian parameter inference for stochastic biochemical network models using particle Markov chain Monte Carlo. Interface Focus 1, 807–820. (doi:10.1098/rsfs.2011.0047).

[103] Warne DJ, Baker RE, Simpson MJ. 2020 A practical guide to pseudo-marginal methods for computational inference in systems biology. Journal of Theoretical Biology 496, 110255. (doi:10.1016/j.jtbi.2020.110255).

[104] Nestel PJ, Whyte HM, Goodman DS. 1969 Distribution and turnover of cholesterol in humans. Journal of Clinical Investigation 48, 982–991. (doi:10.1172/jci106079).

[105] Kermack WO, McKendrick AG. 1927 A contribution to the mathematical theory of epidemics. Proceedings of the Royal Society of London Series A 115, 700–721. (doi:10.1098/rspa.1927.0118).

[106] Tuncer N, Le TT. 2018 Structural and practical identifiability analysis of outbreak models. Mathematical Biosciences 299, 1–18. (doi:10.1016/j.mbs.2018.02.004).

[107] Alahmadi A, Belet S, Black A, Cromer D, Flegg JA, House T, Jayasundara P, Keith JM, McCaw JM, Moss R, Ross JV, Shearer FM, Tun STT, Walker J, White L, Whyte JM, Ada WC, Zarebski AE. 2020 Influencing public health policy with data-informed mathematical models of infectious diseases: Recent developments and new challenges. Epidemics p. 100393. (doi:10.1016/j.epidem.2020.100393).

[108] Topp B, Promislow K, Devries G, Miura RM, Finegood DT. 2000 A model of β-cell mass, insulin, and glucose kinetics: pathways to diabetes. Journal of Theoretical Biology 206, 605–619. (doi:10.1006/jtbi.2000.2150).

[109] Karin O, Swisa A, Glaser B, Dor Y, Alon U. 2016 Dynamical compensation in physiological circuits. Molecular Systems Biology 12, 886. (doi:10.15252/msb.20167216).

[110] Villaverde AF, Tsiantis N, Banga JR. 2019 Full observability and estimation of unknown inputs, states and parameters of nonlinear biological models. Journal of The Royal Society Interface 16, 20190043. (doi:10.1098/rsif.2019.0043).

[111] Engblom S. 2006 Computing the moments of high dimensional solutions of the master equation. Applied Mathematics and Computation 180, 498–515. (doi:10.1016/j.amc.2005.12.032).

[112] Lakatos E, Ale A, Kirk PDW, Stumpf MPH. 2015 Multivariate moment closure techniques for stochastic kinetic models. The Journal of Chemical Physics 143, 094107. (doi:10.1063/1.4929837).

[113] Kuehn C. 2016 Moment closure - a brief review. In Control of Self-Organizing Nonlinear Systems pp. 253–271. (doi:10.1007/978-3-319-28028-8).

[114] Fan S, Geissmann Q, Lakatos E, Lukauskas S, Ale A, Babtie AC, Kirk PDW, Stumpf MPH. 2016 MEANS: python package for Moment Expansion Approximation, iNference and Simulation. Bioinformatics 32, 2863–2865. (doi:10.1093/bioinformatics/btw229).

[115] Brouwer AF, Meza R, Eisenberg MC. 2017 A systematic approach to determining the identifiability of multistage carcinogenesis models. Risk Analysis 37, 1375–1387. (doi:10.1111/risa.12684).

[116] Chiş O, Banga JR, Balsa-Canto E. 2011 GenSSI: a software toolbox for structural identifiability analysis of biological models. Bioinformatics 27, 2610–2611. (doi:10.1093/bioinformatics/btr431).

[117] Ligon TS, Fröhlich F, Chiş OT, Banga JR, Balsa-Canto E, Hasenauer J. 2017 GenSSI 2.0: multi-experiment structural identifiability analysis of SBML models. Bioinformatics 34, 1421–1423. (doi:10.1093/bioinformatics/btx735).

[118] Bezanson J, Edelman A, Karpinski S, Shah VB. 2017 Julia: a fresh approach to numerical computing. SIAM Review 59, 65–98. (doi:10.1137/141000671).

[119] Gillespie DT. 2001 Approximate accelerated stochastic simulation of chemically reacting systems. The Journal of Chemical Physics 115, 1716–1733. (doi:10.1063/1.1378322).

[120] Schnoerr D, Sanguinetti G, Grima R. 2014 The complex chemical Langevin equation. The Journal of Chemical Physics 141, 024103. (doi:10.1063/1.4885345).

[121] Higham DJ. 2008 Modeling and simulating chemical reactions. SIAM Review 50, 347–368. (doi:10.1137/060666457).

[122] Erban R, Chapman SJ. 2009 Stochastic modelling of reaction–diffusion processes: algorithms for bimolecular reactions. Physical Biology 6, 046001. (doi:10.1088/1478-3975/6/4/046001).

[123] Warne DJ, Baker RE, Simpson MJ. 2019 Simulation and inference algorithms for stochastic biochemical reaction networks: from basic concepts to state-of-the-art. Journal of The Royal Society Interface 16, 20180943–20. (doi:10.1098/rsif.2018.0943).

[124] Gillespie DT. 1977 Exact stochastic simulation of coupled chemical reactions. The Journal of Physical Chemistry 81, 2340–2361. (doi:10.1021/j100540a008).

[125] Kurtz TG. 1972 The relationship between stochastic and deterministic models for chemical reactions. The Journal of Chemical Physics 57, 2976–2978. (doi:10.1063/1.1678692).

[126] Gibson MA, Bruck J. 2000 Efficient exact stochastic simulation of chemical systems with many species and many channels. The Journal of Physical Chemistry A 104, 1876–1889. (doi:10.1021/jp993732q).

[127] Rao CV, Wolf DM, Arkin AP. 2002 Control, exploitation and tolerance of intracellular noise. Nature 420, 231–237. (doi:10.1038/nature01258).

[128] Samad HE, Khammash M, Petzold L, Gillespie D. 2005 Stochastic modelling of gene regulatory networks. International Journal of Robust and Nonlinear Control 15, 691–711. (doi:10.1002/rnc.1018).

[129] Golightly A, Wilkinson DJ. 2005 Bayesian inference for stochastic kinetic models using a diffusion approximation. Biometrics 61, 781–788. (doi:10.1111/j.1541-0420.2005.00345.x).

[130] Maruyama G. 1955 Continuous Markov processes and stochastic equations. Rendiconti del Circolo Matematico di Palermo 4, 48. (doi:10.1007/bf02846028).

[131] Socha L. 2008 Linearization Methods for Stochastic Dynamic Systems. Lecture Notes in Physics. Berlin Heidelberg: Springer-Verlag. (doi:10.1007/978-3-540-72997-6).

[132] Hausken K, Moxnes JF. 2010 A closure approximation technique for epidemic models. Mathematical and Computer Modelling of Dynamical Systems 16, 555–574. (doi:10.1080/13873954.2010.496149).

[133] Isserlis L. 1918 On a formula for the product-moment coefficient of any order of a normal frequency distribution in any number of variables. Biometrika 12, 134. (doi:10.2307/2331932).

[134] Singh A, Hespanha JP. 2007 A derivative matching approach to moment closure for the stochastic logistic model. Bulletin of Mathematical Biology 69, 1909–1925. (doi:10.1007/s11538-007-9198-9).

[135] Gelman A, Carlin JB, Stern HS, Dunson DB, Vehtari A, Rubin DB. 2014 Bayesian Data Analysis. CRC Press 3 edition. (doi:10.1201/9780429258411).

[136] Metropolis N, Rosenbluth AW, Rosenbluth MN, Teller AH, Teller E. 1953 Equation of state calculations by fast computing machines. The Journal of Chemical Physics 21, 1087–1092. (doi:10.1063/1.1699114).

[137] Hastings WK. 1970 Monte Carlo sampling methods using Markov chains and their applications. Biometrika 57, 97–109. (doi:10.1093/biomet/57.1.97).

[138] Geyer CJ. 1992 Practical Markov chain Monte Carlo. Statistical Science 7, 473–483. (doi:10.1214/ss/1177011137).

[139] Roberts GO, Rosenthal JS. 2001 Optimal scaling for various Metropolis-Hastings algorithms. Statistical Science 16, 351–367. (doi:10.1214/ss/1015346320).

[140] Gelman A, Rubin DB. 1992 Inference from iterative simulation using multiple sequences. Statistical Science 7, 457–472. (doi:10.1214/ss/1177011136).

[141] Brooks SP, Gelman A. 1998 General methods for monitoring convergence of iterative simulations. Journal of Computational and Graphical Statistics 7, 434. (doi:10.2307/1390675).

[142] Johnston ST, Shah ET, Chopin LK, McElwain DLS, Simpson MJ. 2015 Estimating cell diffusivity and cell proliferation rate by interpreting IncuCyte ZOOM™ assay data using the Fisher-Kolmogorov model. BMC Systems Biology 9, 38. (doi:10.1186/s12918-015-0182-y).

[143] Allen LJS. 2011 An Introduction to Stochastic Processes with Applications to Biology. Boca Raton, Florida: Chapman & Hall/CRC Press.

[144] Matsiaka OM, Baker RE, Shah ET, Simpson MJ. 2019 Mechanistic and experimental models of cell migration reveal the importance of cell-to-cell pushing in cell invasion. Biomedical Physics & Engineering Express 5, 045009. (doi:10.1088/2057-1976/ab1b01).

[145] Golightly A, Wilkinson DJ. 2008 Bayesian inference for nonlinear multivariate diffusion models observed with error. Computational Statistics & Data Analysis 52, 1674–1693. (doi:10.1016/j.csda.2007.05.019).

[146] Poovathingal SK, Gunawan R. 2010 Global parameter estimation methods for stochastic biochemical systems. BMC Bioinformatics 11, 414–414. (doi:10.1186/1471-2105-11-414).

[147] Warne DJ, Ebert A, Drovandi C, Mira A, Mengersen K. 2020 Hindsight is 2020 vision: Characterisation of the global response to the COVID-19 pandemic. medRxiv. (doi:10.1101/2020.04.30.20085662).

[148] Rackauckas C, Nie Q. 2016 DifferentialEquations.jl – A performant and feature-rich ecosystem for solving differential equations in Julia. Journal of Open Research Software 5. (doi:10.5334/jors.151).

[149] Burchard H, Deleersnijder E, Meister A. 2003 A high-order conservative Patankar-type discretisation for stiff systems of production–destruction equations. Applied Numerical Mathematics 47, 1–30. (doi:10.1016/s0168-9274(03)00101-6).

[150] Villaverde AF, Banga JR. 2014 Reverse engineering and identification in systems biology: strategies, perspectives and challenges. Journal of The Royal Society Interface 11, 20130505. (doi:10.1098/rsif.2013.0505).

[151] Villaverde AF. 2019 Observability and structural identifiability of nonlinear biological systems. Complexity 2019, 1–12. (doi:10.1155/2019/8497093).

[152] Brown KS, Sethna JP. 2003 Statistical mechanical approaches to models with many poorly known parameters. Physical Review E 68, 021904. (doi:10.1103/physreve.68.021904).

[153] Transtrum MK, Machta BB, Brown KS, Daniels BC, Myers CR, Sethna JP. 2015 Perspective: Sloppiness and emergent theories in physics, biology, and beyond. The Journal of Chemical Physics 143, 010901. (doi:10.1063/1.4923066).

[154] Dufresne E, Harrington HA, Raman DV. 2018 The geometry of sloppiness. Journal of Algebraic Statistics 9, 30–68. (doi:10.18409/jas.v9i1.64).

[155] Smadbeck P, Kaznessis YN. 2013 A closure scheme for chemical master equations. Proceedings of the National Academy of Sciences 110, 14261–14265. (doi:10.1073/pnas.1306481110).

[156] Zechner C, Ruess J, Krenn P, Pelet S, Peter M, Lygeros J, Koeppl H. 2012 Moment-based inference predicts bimodality in transient gene expression. Proceedings of the National Academy of Sciences 109, 8340–8345. (doi:10.1073/pnas.1200161109).

[157] Roberts GO, Rosenthal JS. 2009 Examples of adaptive MCMC. Journal of Computational and Graphical Statistics 18, 349–367. (doi:10.1198/jcgs.2009.06134).

[158] Moral PD, Doucet A, Jasra A. 2006 Sequential Monte Carlo samplers. Journal of the Royal Statistical Society: Series B 68, 411–436. (doi:10.1111/j.1467-9868.2006.00553.x).

[159] Giles MB. 2008 Multilevel Monte Carlo path simulation. Operations Research 56, 607–617. (doi:10.1287/opre.1070.0496).

[160] Jasra A, Kamatani K, Law KJH, Zhou Y. 2017 Multilevel particle filters. SIAM Journal on Numerical Analysis 55, 3068–3096. (doi:10.1137/17m1111553).

[161] Warne DJ, Baker RE, Simpson MJ. 2018 Multilevel rejection sampling for approximate Bayesian computation. Computational Statistics & Data Analysis 124, 71–86. (doi:10.1016/j.csda.2018.02.009).

[162] Quiroz M, Kohn R, Villani M, Tran MN. 2019 Speeding up MCMC by efficient data subsampling.. Journal of the American Statistical Association 114, 831–843. (doi:10.1080/01621459.2018.1448827).

[163] Pooley CM, Bishop SC, Marion G. 2015 Using model-based proposals for fast parameter inference on discrete state space, continuous-time Markov processes. Journal of The Royal Society Interface 12, 20150225. (doi:10.1098/rsif.2015.0225).

[164] Burrage K, Burrage P. 1996 High strong order explicit Runge-Kutta methods for stochastic ordinary differential equations. Applied Numerical Mathematics 22, 81–101. (doi:10.1016/S0168-9274(96)00027-X).

[165] Mingas G, Bottolo L, Bouganis CS. 2017 Particle MCMC algorithms and architectures for accelerating inference in state-space models. International Journal of Approximate Reasoning 83, 413–433. (doi:10.1016/j.ijar.2016.10.011).

[166] Lee YS, Liu OZ, Hwang HS, Knollmann BC, Sobie EA. 2013 Parameter sensitivity analysis of stochastic models provides insights into cardiac calcium sparks. Biophysical Journal 104, 1142–1150. (doi:10.1016/j.bpj.2012.12.055).

[167] Botha I, Kohn R, Drovandi C. 2020 Particle methods for stochastic differential equation mixed effects models. Bayesian Analysis. (doi:10.1214/20-ba1216).

[168] Picchini U. 2012 Inference for SDE models via approximate Bayesian computation. Journal of Computational and Graphical Statistics 23, 1080–1100. (doi:10.1080/10618600.2013.866048).

[169] Buckwar E, Tamborrino M, Tubikanec I. 2020 Spectral density-based and measure-preserving ABC for partially observed diffusion processes. An illustration on Hamiltonian SDEs. Statistics and Computing 30, 627–648. (doi:10.1007/s11222-019-09909-6).

[170] Liepe J, Filippi S, Komorowski M, Stumpf MPH. 2013 Maximizing the information content of experiments in systems biology. PLOS Computational Biology 9, e1002888. (doi:10.1371/journal.pcbi.1002888).

[171] Evans ND, White LJ, Chapman MJ, Godfrey KR, Chappell MJ. 2005 The structural identifiability of the susceptible infected recovered model with seasonal forcing. Mathematical Biosciences 194, 175–197. (doi:10.1016/j.mbs.2004.10.011).

[172] Chapman JD, Evans ND. 2008 The structural identifiability of SIR type epidemic models with incomplete immunity and birth targeted vaccination. IFAC Proceedings Volumes 41, 9075–9080. (doi:10.3182/20080706-5-kr-1001.01532).

[173] Beskos A, Papaspiliopoulos O, Roberts GO. 2006 Retrospective exact simulation of diffusion sample paths with applications. Bernoulli 12, 1077–1098. (doi:10.3150/bj/1165269151).

[174] Fröhlich F, Thomas P, Kazeroonian A, Theis FJ, Grima R, Hasenauer J. 2016 Inference for stochastic chemical kinetics using moment equations and system size expansion. PLOS Computational Biology 12, e1005030. (doi:10.1371/journal.pcbi.1005030).

[175] Plotnikov A, Zehorai E, Procaccia S, Seger R. 2011 The MAPK cascades: Signaling components, nuclear roles and mechanisms of nuclear translocation. Biochimica et Biophysica Acta (BBA) - Molecular Cell Research 1813, 1619–1633. (doi:10.1016/j.bbamcr.2010.12.012).

[176] Xiu D, Karniadakis GE. 2002 The Wiener-Askey polynomial chaos for stochastic differential equations. SIAM Journal on Scientific Computing 24, 619–644. (doi:10.1137/s1064827501387826).

[177] Archambeau C, Cornford D, Opper M, Shawe-Taylor JS. 2007 Gaussian process approximations of stochastic differential equations. Proceedings of Machine Learning Research 1, 1–16.

[178] Mirams GR, Pathmanathan P, Gray RA, Challenor P, Clayton RH. 2016 Uncertainty and variability in computational and mathematical models of cardiac physiology. The Journal of Physiology 594, 6833–6847. (doi:10.1113/jp271671).

[179] Kaintura A, Dhaene T, Spina D. 2018 Review of polynomial chaos-based methods for uncertainty quantification in modern integrated circuits. Electronics 7, 30. (doi:10.3390/electronics7030030).

[180] Ran ZY, Hu BG. 2017 Parameter identifiability in statistical machine learning: a review. Neural Computation 29, 1151–1203. (doi:doi.org/10.1162/neco_a_00947).

[181] Lill D, Timmer J, Kaschek D. 2019 Local Riemannian geometry of model manifolds and its implications for practical parameter identifiability. PLOS ONE 14, e0217837. (doi:10.1371/journal.pone.0217837).

[182] Livingstone S, Girolami M. 2014 Information-geometric Markov chain Monte Carlo methods using diffusions. Entropy 16, 3074–3102. (doi:10.3390/e16063074).

[183] Archambeau C, Opper M, Shen Y, Cornford D, Shawe-Taylor JS. 2008 Variational inference for diffusion processes. In Platt JC, Koller D, Singer Y, Roweis ST, editors, Advances in Neural Information Processing Systems vol. 20 pp. 17–24. Curran Associates, Inc.

[184] Raue A, Kreutz C, Theis FJ, Timmer J. 2013 Joining forces of Bayesian and frequentist methodology: a study for inference in the presence of non-identifiability. Philosophical Transactions of the Royal Society A: Mathematical, Physical and Engineering Sciences 371, 20110544. (doi:10.1098/rsta.2011.0544).

[185] Faller D, Klingmüller U, Timmer J. 2003 Simulation methods for optimal experimental design in systems biology. SIMULATION 79, 717–725. (doi:10.1177/0037549703040937).

[186] Walter E, Lecourtier Y. 1981 Unidentifiable compartmental models: what to do?. Mathematical Biosciences 56, 1–25. (doi:10.1016/0025-5564(81)90025-0).

