## Supporting Material Document for "Identifiability analysis for stochastic differential equation models in systems biology"

#### Code availability

This supporting material document is supplementary to, and refers to, code available on Github at github.com/ap-browning/SDE-Identifiability.

### 1 Stochastic simulation algorithm

We simulate the first three models as bio-chemical reaction networks with an event-driven stochastic simulation algorithm (SSA), developed by Gillespie [1]. Each model comprises $Q$ reactions, with propensities and stoichiometries denoted

$$\{(a_k(\mathbf{X}_t; \boldsymbol{\theta}), \boldsymbol{\nu}_k)\}_{k=1}^{Q}, \tag{1}$$

respectively. We simulate from the initial condition $\mathbf{X}_0$ at $t = 0$ to the final state $\mathbf{X}_{t_f}$ at $t = t_f$. While simulating from the SSA, $\mathbf{X}_t \in \mathbb{Z}^N$. We summarise the SSA in algorithm 1 and provide our implementation in `Module/Models/SimulateSSA.jl`.

**Algorithm 1** Stochastic simulation algorithm

1: Initialise by setting $t = 0$ and $\mathbf{X}_t = \mathbf{X}_0$.
2: Compute total event rate, $a(\mathbf{X}_t) = \sum_{k=1}^{Q} a_k(\mathbf{X}_t; \boldsymbol{\theta})$.
3: Sample time-step $\Delta t \sim \text{Exp}(a(\mathbf{X}_t))$. If $t + \Delta t > t_f$ stop the algorithm. Else, set $t \leftarrow t + \Delta t$.
4: Sample event, $k$, such that $\mathbb{P}(k) = a_k(\mathbf{X}_t; \boldsymbol{\theta})/a(\mathbf{X}_t)$.
5: Update state, $\mathbf{X}_t \leftarrow \mathbf{X}_t + \boldsymbol{\nu}_k$.
6: Repeat steps 2 – 6.

### 2 Euler-Maruyama algorithm

We simulate Itô SDEs using the Euler-Maruyama algorithm [2] with reflecting boundaries at $X_{i,t} = 0$ to ensure positivity [3]. Each SDE is defined by

$$\mathrm{d}\mathbf{X}_t = \boldsymbol{\alpha}(\mathbf{X}_t, t; \boldsymbol{\theta})\,\mathrm{d}t + \boldsymbol{\sigma}(\mathbf{X}_t, t; \boldsymbol{\theta})\,\mathrm{d}\mathbf{W}_t. \tag{2}$$

Here $\mathbf{X}_t$ is an $N$-dimensional vector; $\boldsymbol{\alpha}(\cdot)$ maps to an $N$-dimensional vector; $\boldsymbol{\sigma}(\cdot)$ maps to an $N \times Q$ matrix; and $\mathbf{W}_t$ is an $Q$-dimensional Wiener process with independent components. We simulate from the initial condition $\mathbf{X}_{t_0}$ at $t = t_0$ to the final state $\mathbf{X}_{t_f}$ at $t = t_f$. We summarise the Euler-Maruyama algorithm in algorithm 2 and provide our implementation in `Module/Models/EulerMaruyama.jl`.

**Algorithm 2** Euler-Maruyama algorithm

1: Initialise by setting $t = t_0$ and $\mathbf{X}_t = \mathbf{X}_{t_0}$. Choose time-step, $\Delta t$.
2: Sample Wiener increment $\Delta\mathbf{W} = (\Delta W_1, \Delta W_2, \ldots, \Delta W_Q)^T$ where $\Delta W_k \sim \mathcal{N}(0, \Delta t)$.
3: Update state $\mathbf{X}_t \leftarrow \mathbf{X}_t + \boldsymbol{\alpha}(\mathbf{X}_t, t; \boldsymbol{\theta})\Delta t + \boldsymbol{\sigma}(\mathbf{X}_t, t; \boldsymbol{\theta})\Delta\mathbf{W}$.
4: Set $X_{i,t} = |X_{i,t}|$ where $\mathbf{X}_t = (X_{1,t}, X_{2,t}, \ldots, X_{N,t})^T$.
5: Update time, $t \leftarrow t + \Delta t$.
6: Repeat steps 2 – 5 until $t \geq t_f$.

### 3 MCMC algorithms

#### 3.1 Metropolis-Hastings algorithm

We simulate from the posterior distribution

$$p(\boldsymbol{\theta}|\mathcal{D}) \propto \mathcal{L}(\mathcal{D}|\boldsymbol{\theta})p(\boldsymbol{\theta}), \tag{3}$$

using the Metropolis-Hastings algorithm [4, 5] and Markov-chain Monte-Carlo (MCMC). In this work, we employ a multivariate normal proposal kernel with covariance $\boldsymbol{\Sigma}$ such that

$$q(\boldsymbol{\theta}|\boldsymbol{\theta}_{s-1}) = \text{MVN}(\boldsymbol{\theta}_{s-1}, \boldsymbol{\Sigma}). \tag{4}$$

We summarise our implementation of the Metropolis-Hastings MCMC algorithm in algorithm 3 and provide our implementation in `Module/Inference/MetropolisHastings.jl`.

**Algorithm 3** Metropolis-Hastings algorithm with a symmetric proposal kernel.

1: Sample $\boldsymbol{\theta}_0 \sim p(\boldsymbol{\theta})$ to initialise chain at $s = 0$. Choose desired number of iterations, $S$.
2: Update iteration $s \leftarrow s + 1$.
3: Propose transition $\boldsymbol{\theta}^* \sim q(\boldsymbol{\theta}|\boldsymbol{\theta}_{s-1})$, where $q(\boldsymbol{\theta}|\boldsymbol{\theta}_{s-1})$ is a multivariate Gaussian.
4: Calculate acceptance probability

$$\alpha_{\text{MH}}(\boldsymbol{\theta}^*|\boldsymbol{\theta}_{s-1}) = \min\left(1, \frac{p(\boldsymbol{\theta}^*)\mathcal{L}(\mathcal{D}|\boldsymbol{\theta}^*)}{p(\boldsymbol{\theta}_{s-1})\mathcal{L}(\mathcal{D}|\boldsymbol{\theta}_{s-1})}\right).$$

5: Accept proposal, $\boldsymbol{\theta}_s \leftarrow \boldsymbol{\theta}^*$ with probability $\alpha_{\text{MH}}(\boldsymbol{\theta}^*|\boldsymbol{\theta}_{s-1})$; else, reject and set $\boldsymbol{\theta}_s \leftarrow \boldsymbol{\theta}_{s-1}$.
6: Repeat steps 2 – 5 until $s = S$.

#### 3.2 Particle MCMC algorithm

For most SDE models, the likelihood function, $\mathcal{L}(\mathcal{D}|\boldsymbol{\theta})$, is intractable. We approximate the likelihood using a bootstrap particle filter in a particle MCMC algorithm [6], replacing $\mathcal{L} \leftarrow \hat{\mathcal{L}}$ in algorithm 3.

The data are defined by $\mathcal{D} = \left\{\{t_{n,i}, \mathbf{Y}_{\text{obs}}^{n,i}\}_{n=1}^{N_E}\right\}_{i=1}^{E}$, where

$$\mathbf{Y}_{\text{obs}}^{n,i} \sim g(\mathbf{Y}|\mathbf{X}_{t_n}^i, t_n; \boldsymbol{\theta}). \tag{5}$$

Here, $\{\mathbf{X}_{t_n}^i\}_{n=1}^{N_E}$ are observations from experiment $i = 1, 2, \ldots, N_E$, modelled by a single trace of the SDE (equation (2)) initiated at $\mathbf{X}_0 = \mathbf{h}_i(\boldsymbol{\theta})$. The vector-valued function $\mathbf{h}_i(\boldsymbol{\theta})$ captures the fact that the initial condition may depend on unknown parameters, $\boldsymbol{\theta}$. This occurs in the epidemic model where only partial observations of the state are made so that the full initial condition is unknown.

We summarise our implementation of bootstrap particle filter algorithm in algorithm 4 and provide our implementation in `Module/Inference/PMLogLikeParticleFilter.jl`.

**Algorithm 4** Bootstrap particle filter

1: Initialise likelihood estimate, $\hat{\mathcal{L}} = 1$, and experiment index $i = 0$.

2: Update experiment index $i \leftarrow i+1$ and initialise $R$ particles for the $i$th experiment, $\{\mathbf{X}_{t_0}^r\}_{r=1}^R$, where $\mathbf{X}_0^r = \mathbf{h}_i(\boldsymbol{\theta})$ and $t_0 = 0$. Set observation index, $n = 0$.

3: Update observation index, $n \leftarrow n+1$, and simulate all particles forward from $t = t_{n-1}$ to $t_n$ using the Euler-Maruyama algorithm (algorithm 2) to obtain $\{\mathbf{X}_{t_n}^r\}_{r=1}^R$.

4: Compute weight of each particle, $\{W_n^r\}$, such that $W_n^r = g(\mathbf{Y}_{\text{obs}}^{n,i}|\mathbf{X}_{t_n}^r)$.

5: Update likelihood estimate,

$$\hat{\mathcal{L}} \leftarrow \hat{\mathcal{L}} \times \frac{1}{R}\sum_{r=1}^{R} W_n^r.$$

6: Resample $R$ particles from $\{\mathbf{X}_{t_n}^r\}_{r=1}^R$ with replacement according to weights $\{W_n^r\}$.

7: Repeat steps 3 – 6 until $n = N_E$.

8: Repeat steps 2 – 7 until $i = E$.

#### 3.3 Likelihood for ODE models

When describing data using an ODE model, we make the typical assumption that residuals are independent. The likelihood function is then tractable and given by

$$\mathcal{L} = \sum_{i=1}^{E}\sum_{n=1}^{N_E} g\big(\mathbf{Y}_{\text{obs}}^{n,i}|\mathbf{X}_{t_n}^i, t_n; \boldsymbol{\theta}\big). \tag{6}$$

Here, $\{\mathbf{X}_{t_n}^i\}_{n=1}^{N_E}$ are observations from experiment $i$ ($i = 1, 2, \ldots, N_E$), modelled by the ODE model initiated at $\mathbf{X}_0 = \mathbf{h}_i(\boldsymbol{\theta})$.

#### 4 Moment equations

In the main text, we define the raw moments,

$$m_{i_1 i_2 \dots i_N}(t) = \left\langle \prod_{j=1}^{N} X_{j,t}^{i_j} \right\rangle, \tag{7}$$

of the random variable $\mathbf{X}_t$, which is itself described by the Itô SDE (equation (2)). Here, we derive the equation describing the time-evolution of the moments.

First, we define $\phi_{\mathbf{i}}(\mathbf{X}_t, t) = \prod_{j=1}^{N} X_{j,t}^{i_j}$, where $\mathbf{i} = (i_1, i_2, \dots, i_N)$. By Itô's lemma,

$$\mathrm{d}\phi_{\mathbf{i}}(\mathbf{X}, t) = f_\phi(\mathbf{X}_t, t; \boldsymbol{\theta})\mathrm{d}t + \sum_{i=1}^{N} \mathbf{g}_\phi(\mathbf{X}_t, t; \boldsymbol{\theta})\mathrm{d}\mathbf{W}_t, \tag{8}$$

where

$$f_\phi(\mathbf{X}_t, t; \boldsymbol{\theta}) = \boldsymbol{\alpha}(\mathbf{X}_t, t; \boldsymbol{\theta}) \cdot \boldsymbol{\nabla} \left( \prod_{j=1}^{N} X_{j,t}^{i_j} \right) + \frac{1}{2}\mathrm{Tr}\left( \boldsymbol{\sigma}^T(\mathbf{X}_t, t; \boldsymbol{\theta}) \mathbf{H} \left( \prod_{j=1}^{N} X_j^{i_j} \right) \boldsymbol{\sigma}(\mathbf{X}_t, t; \boldsymbol{\theta}) \right), \tag{9}$$

$$\mathbf{g}_\phi(\mathbf{X}_t, t; \boldsymbol{\theta}) = \boldsymbol{\nabla} \left( \prod_{j=1}^{N} X_{j,t}^{i_j} \right)^T \boldsymbol{\sigma}(\mathbf{X}_t, t; \boldsymbol{\theta}). \tag{10}$$

Our aim is to find an ODE describing the time evolution of $\langle \phi_{\mathbf{i}}(\mathbf{X}) \rangle$, where $\langle \cdot \rangle$ is the expectation taken with respect to the probability measure of the random variable $\mathbf{X}_t$. Full details are available in [7].

Consider the expectation of equation (8) in integral form:

$$\langle \phi_{\mathbf{i}}(\mathbf{X}, t) \rangle = \langle \phi_{\mathbf{i}}(\mathbf{X}, 0) \rangle + \underbrace{\left\langle \int_0^t f_\phi(\mathbf{X}_u, u; \boldsymbol{\theta})\mathrm{d}u \right\rangle}_{(*)} + \underbrace{\sum_{n=1}^{N} \left\langle \int_0^t g_\phi^{(n)}(\mathbf{X}_u, u; \boldsymbol{\theta})\mathrm{d}W_{n,u} \right\rangle}_{(**)}. \tag{11}$$

Here, we denote $\mathbf{W} = (W_{1,t}, W_{2,t} \dots, W_{N,t})^T$ and $\mathbf{g}_\phi(\cdot) = \left( g_\phi^{(1)}(\cdot), g_\phi^{(2)}(\cdot), \dots, g_\phi^{(N)}(\cdot) \right)^T$. Under certain conditions, which are satisfied when all elements of $\mathbf{g}_\phi$ are polynomial, the stochastic integrals in equation (11)(∗∗) will vanish [8] due to the Itô formulation. This is the case for the SDEs we consider, which are derived through the chemical Langevin equation with polynomial propensity functions, so that all components of $\mathbf{g}_\phi$ are also polynomials. Therefore, after interchanging the order of time-integration to that of the expectation in (∗), we obtain

$$\langle \phi_{\mathbf{i}}(\mathbf{X}, t) \rangle = \langle \phi_{\mathbf{i}}(\mathbf{X}, 0) \rangle + \int_0^t \langle f_\phi(\mathbf{X}_u, u; \boldsymbol{\theta}) \rangle \,\mathrm{d}u. \tag{12}$$

We note that, by definition, $m_{i_1 i_2 \dots i_N}(t) = \langle \phi_{\mathbf{i}}(\mathbf{X}, t) \rangle$, and so

$$m_{i_1 i_2 \dots i_N}(t) = m_{i_1 i_2 \dots i_N}(0) + \int_0^t \langle f_\phi(\mathbf{X}_u, u; \boldsymbol{\theta}) \rangle \,\mathrm{d}u, \tag{13}$$

which corresponds to the ODE

$$\frac{\mathrm{d}m_{i_1 i_2 \dots i_N}}{\mathrm{d}t} = \langle f_\phi(\mathbf{X}_u, u; \boldsymbol{\theta}) \rangle . \tag{14}$$

We provide `Mathematica` scripts used to derive the moment equations for each model in the `Mathematica` folder on Github.

#### 4.1 Epidemic model

The CLE for the epidemic model is

$$\mathrm{d}\mathbf{X}_t = \begin{pmatrix} -\theta_1 X_{1,t} X_{3,t} \\ \theta_1 X_{1,t} X_{3,t} - \theta_2 X_{2,t} \\ \theta_2 X_{2,t} - \theta_3 X_{3,t} \\ \theta_3 X_{3,t} \end{pmatrix} \mathrm{d}t + \begin{pmatrix} -\sqrt{\theta_1 X_{1,t} X_{3,t}} & 0 & 0 \\ \sqrt{\theta_1 X_{1,t} X_{3,t}} & -\sqrt{\theta_2 X_{2,t}} & 0 \\ 0 & \sqrt{\theta_2 X_{2,t}} & -\sqrt{\theta_3 X_{3,t}} \\ 0 & 0 & \sqrt{\theta_3 X_{3,t}} \end{pmatrix} \mathrm{d}\mathbf{W}_t. \tag{15}$$

We denote

$$m_{i_1 i_2 i_3 i_4}(t) = \left\langle X_{1,t}^{i_1} X_{2,t}^{i_2} X_{3,t}^{i_3} X_{4,t}^{i_4} \right\rangle . \tag{16}$$

To second order, the unclosed moment equations are

$$\left.\begin{aligned}
\frac{\mathrm{d}m_{1000}}{\mathrm{d}t} &= -\theta_1 m_{1010}, \\
\frac{\mathrm{d}m_{0100}}{\mathrm{d}t} &= \theta_1 m_{1010} - \theta_2 m_{0100}, \\
\frac{\mathrm{d}m_{0010}}{\mathrm{d}t} &= \theta_2 m_{0100} - \theta_3 m_{0010}, \\
\frac{\mathrm{d}m_{0001}}{\mathrm{d}t} &= \theta_3 m_{0010}, \\
\frac{\mathrm{d}m_{2000}}{\mathrm{d}t} &= -2\theta_1 m_{2010} + \theta_1 m_{1010}, \\
\frac{\mathrm{d}m_{0200}}{\mathrm{d}t} &= 2\theta_1 m_{1110} + \theta_1 m_{1010} + \theta_2 m_{0100} - 2\theta_2 m_{0200}, \\
\frac{\mathrm{d}m_{0020}}{\mathrm{d}t} &= \theta_2 m_{0100} + 2\theta_2 m_{0110} + \theta_3 m_{0010} - 2\theta_3 m_{0020}, \\
\frac{\mathrm{d}m_{0002}}{\mathrm{d}t} &= \theta_3 m_{0010} + 2\theta_3 m_{0011}, \\
\frac{\mathrm{d}m_{1100}}{\mathrm{d}t} &= -\theta_1 m_{1110} + \theta_1 m_{2010} - \theta_1 m_{1010} - \theta_2 m_{1100}, \\
\frac{\mathrm{d}m_{1010}}{\mathrm{d}t} &= -\theta_1 m_{1020} + \theta_2 m_{1100} - \theta_3 m_{1010}, \\
\frac{\mathrm{d}m_{1001}}{\mathrm{d}t} &= -\theta_1 m_{1011} + \theta_3 m_{1010}, \\
\frac{\mathrm{d}m_{0110}}{\mathrm{d}t} &= \theta_1 m_{1020} - \theta_2 m_{0100} - \theta_2 m_{0110} + \theta_2 m_{0200} - \theta_3 m_{0110}, \\
\frac{\mathrm{d}m_{0101}}{\mathrm{d}t} &= \theta_1 m_{1011} - \theta_2 m_{0101} + \theta_3 m_{0110}, \\
\frac{\mathrm{d}m_{0011}}{\mathrm{d}t} &= \theta_2 m_{0101} - \theta_3 m_{0010} - \theta_3 m_{0011} + \theta_3 m_{0020}.
\end{aligned}\right\} \tag{17}$$

Here, we have indicated moments of order three and above in red font. To close the system, we approximate the higher order moments using the following closures.

*Mean-field.* The mean-field closure approximates third order moments with the appropriate product of first order moments. This results in the following approximations for each of the third-order raw moments in equation (17):

$$m_{2010} \approx m_{1000}^2 m_{0010}, \tag{18a}$$
$$m_{1110} \approx m_{1000} m_{0100} m_{0010}, \tag{18b}$$
$$m_{1020} \approx m_{1000} m_{0020}^2, \tag{18c}$$
$$m_{1011} \approx m_{1000} m_{0010} m_{0001}. \tag{18d}$$

*Pair-wise.* The pair-wise closure approximates third order moments with a product and quotient of first and second order moments. This results in the following approximations for each of the third-order raw moments in equation (17):

$$m_{2010} \approx \frac{m_{2000} m_{1010}}{m_{1000}}, \tag{19a}$$
$$m_{1110} \approx \frac{m_{1100} m_{0110}}{m_{0100}}, \tag{19b}$$
$$m_{1020} \approx \frac{m_{1010} m_{0020}}{m_{0010}}, \tag{19c}$$
$$m_{1011} \approx \frac{m_{1010} m_{0011}}{m_{0010}}. \tag{19d}$$

*Gaussian.* The Gaussian closure sets the third order *central* moments to zero. That is,

$$0 = \hat{m}_{i_1 i_2 i_3 i_4}(t) = \left\langle \prod_{j=1}^{4} \left( X_{i,t} - \langle X_{i,t} \rangle \right)^{i_j} \right\rangle.$$

This results in the following approximations for each of the third-order raw moments in equation (17):

$$m_{2010} \approx -2m_{0010} m_{1000}^2 + 2m_{1000} m_{1010} + m_{0010} m_{2000}, \tag{20a}$$
$$m_{1110} \approx -2m_{0010} m_{0100} m_{1000} + m_{0110} m_{1000} + m_{0100} m_{1010} + m_{0010} m_{1100}, \tag{20b}$$
$$m_{1020} \approx -2m_{0010} m_{1000}^2 + 2m_{0010} m_{1010} + m_{0020} m_{1000}, \tag{20c}$$
$$m_{1011} \approx -2m_{0001} m_{0010} m_{1000} + m_{0011} m_{1000} + m_{0010} m_{1001} + m_{0001} m_{1010}. \tag{20d}$$

##### 4.1.1 Comparison of closures

In figure S1 we compare the ODE model to the mean-field closure, pair-wise closure and Gaussian closure for the epidemic model. The mean-field model was solved using the positivity-preserving Patankar-type method [9] (of order one) with time-step $\Delta t = 0.001$, the other closures and the ODE model using the `Tsit5` routine (a 4th and 5th order Runge-Kutte regime) in `Julia` [10].

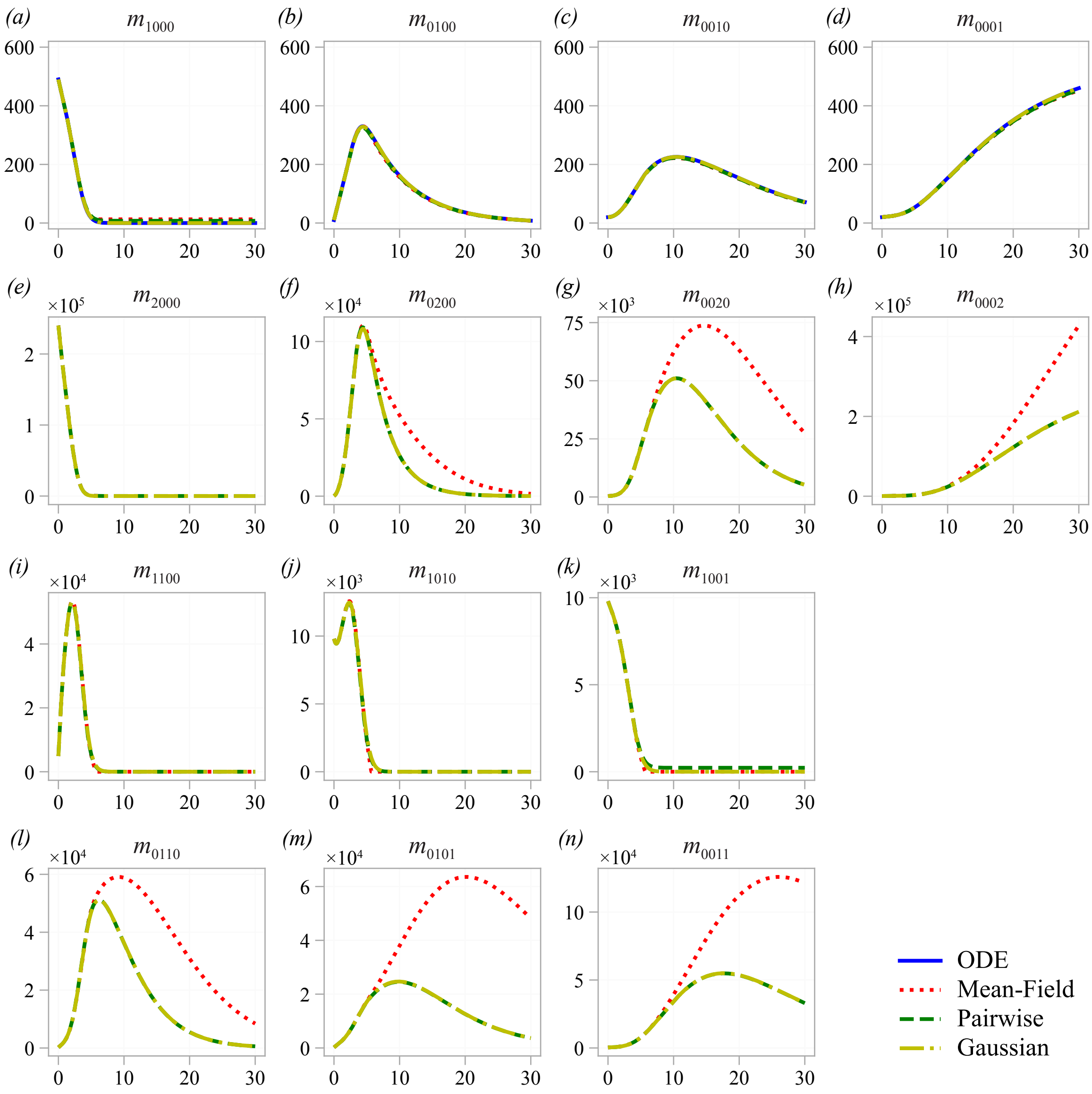

**Figure S1.** Comparison of closure methods for the epidemic model.

#### References

[1] Gillespie DT. 1977 Exact stochastic simulation of coupled chemical reactions. *The Journal of Physical Chemistry* **81**, 2340–2361. (doi:10.1021/j100540a008).

[2] Maruyama G. 1955 Continuous Markov processes and stochastic equations. *Rendiconti del Circolo Matematico di Palermo* **4**, 48. (doi:10.1007/bf02846028).

[3] Dangerfield CE, Kay D, Burrage K. 2012 Modeling ion channel dynamics through reflected stochastic differential equations. *Physical Review E* **85**, 051907. (doi:10.1103/physreve.85.051907).

[4] Metropolis N, Rosenbluth AW, Rosenbluth MN, Teller AH, Teller E. 1953 Equation of state calculations by fast computing machines. *The Journal of Chemical Physics* **21**, 1087–1092. (doi:10.1063/1.1699114).

[5] Hastings WK. 1970 Monte Carlo sampling methods using Markov chains and their applications. *Biometrika* **57**, 97–109. (doi:10.1093/biomet/57.1.97).

[6] Andrieu C, Roberts GO. 2009 The pseudo-marginal approach for efficient Monte Carlo computations. *The Annals of Statistics* **37**, 697–725. (doi:10.1214/07-aos574).

[7] Socha L. 2008 *Linearization Methods for Stochastic Dynamic Systems.* Lecture Notes in Physics. Berlin Heidelberg: Springer-Verlag. (doi:10.1007/978-3-540-72997-6).

[8] Kloeden PE, Platen E. 1992 *Numerical Solution of Stochastic Differential Equations.* Berlin: Springer-Verlag. (doi:10.1007/978-3-662-12616-5).

[9] Burchard H, Deleersnijder E, Meister A. 2003 A high-order conservative Patankar-type discretisation for stiff systems of production–destruction equations. *Applied Numerical Mathematics* **47**, 1–30. (doi:10.1016/s0168-9274(03)00101-6).

[10] Rackauckas C, Nie Q. 2016 DifferentialEquations.jl – A performant and feature-rich ecosystem for solving differential equations in Julia. *Journal of Open Research Software* **5**. (doi:10.5334/jors.151).
